# Topography of mutational signatures in human cancer

**DOI:** 10.1101/2022.05.29.493921

**Authors:** Burcak Otlu, Marcos Díaz-Gay, Ian Vermes, Erik N Bergstrom, Maria Zhivagui, Mark Barnes, Ludmil B. Alexandrov

## Abstract

The somatic mutations found in a cancer genome are imprinted by different mutational processes. Each process exhibits a characteristic mutational signature, which can be affected by the genome architecture. However, the interplay between mutational signatures and topographical genomic features has not been extensively explored. Here, we integrate mutations from 5,120 whole-genome sequenced tumours from 40 cancer types with 516 topographical features from ENCODE to evaluate the effect of nucleosome occupancy, histone modifications, CTCF binding, replication timing, and transcription/replication strand asymmetries on the cancer-specific accumulation of mutations from distinct mutagenic processes. Most mutational signatures are affected by topographical features with signatures of related aetiologies being similarly affected. Certain signatures exhibit periodic behaviours or cancer-type specific enrichments/depletions near topographical features, revealing further information about the processes that imprinted them. Our findings, disseminated via COSMIC, provide a comprehensive online resource for exploring the interactions between mutational signatures and topographical features across human cancer.

**GRAPHICAL ABSTRACT:** 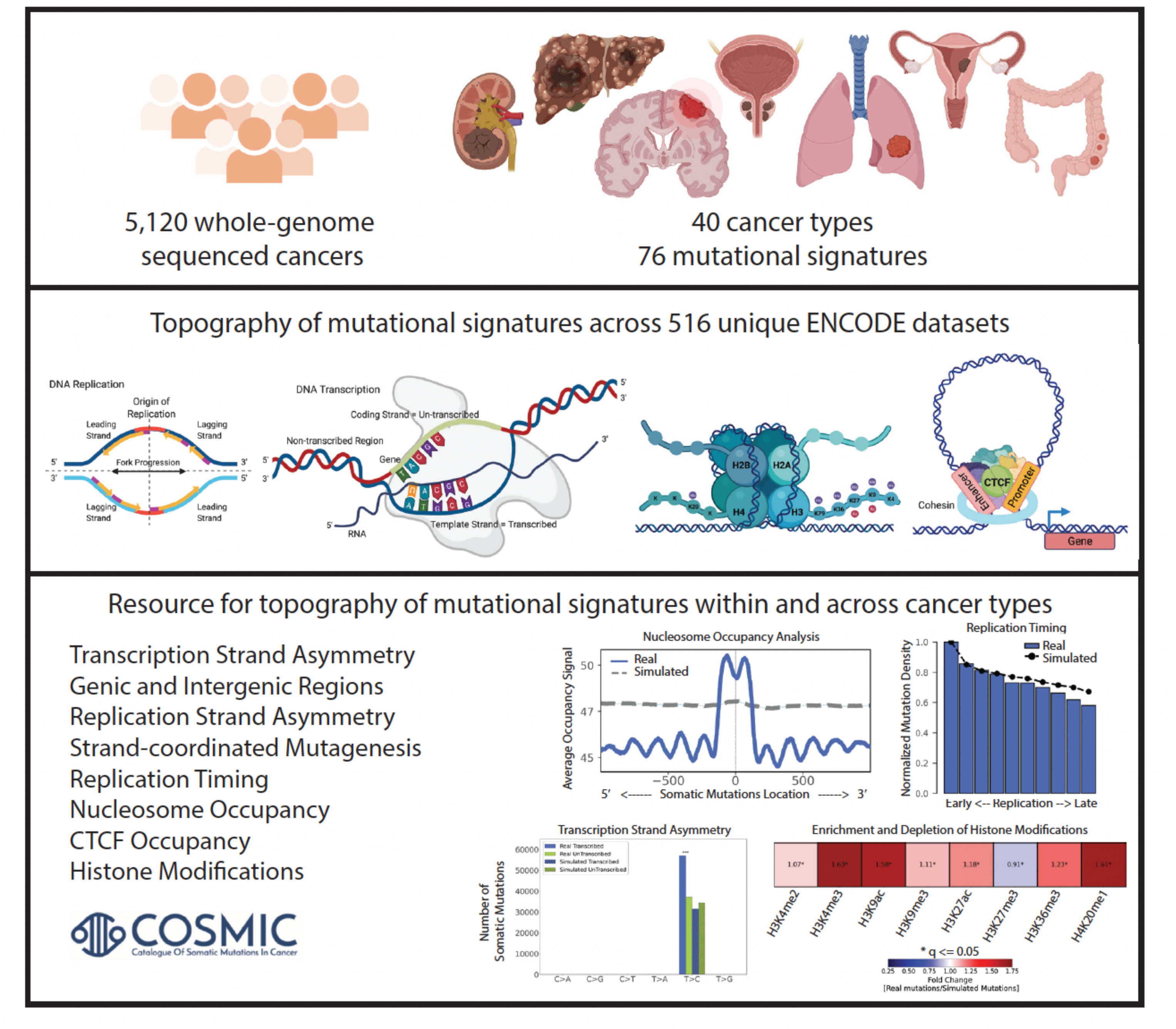

**HIGHLIGHTS:**

- Comprehensive topography analysis of mutational signatures encompassing 82,890,857 somatic mutations in 5,120 whole-genome sequenced tumours integrated with 516 tissue-matched topographical features from the ENCODE project.
- The accumulation of somatic mutations from most mutational signatures is affected by nucleosome occupancy, histone modifications, CTCF binding sites, transcribed regions, or replication strand/timing.
- Mutational signatures with related aetiologies are consistently characterized by similar genome topographies across tissue types.
- Topography analysis allows both separating signatures from different aetiologies and understanding the genomic specificity of clustered somatic mutations.
- A comprehensive online resource, disseminate through the COSMIC signatures database, that allows researchers to explore the interactions between somatic mutational processes and genome architecture within and across cancer types.

## INTRODUCTION

Cancer genomes are peppered with somatic mutations imprinted by the activities of different endogenous and exogenous mutational processes^1, 2^. Due to their intrinsic biophysical and biochemical properties, each mutational process engraves a characteristic pattern of somatic mutations, known as a mutational signature^3^. Our previous analyses encompassing more than 5,000 whole-genome and 20,000 whole-exome sequenced human cancers have revealed the existence of at least 78 single base substitution (SBS), 11 doublet-base substitution (DBS), and 18 indel (ID) mutational signatures^4–7^. Through statistical associations and further experimental characterizations, aetiology has been proposed for approximately half of the identified signatures^4, 8–15^. Prior studies have also explored the interactions between somatic mutations imprinted by different mutational processes and the topographical features of the human genome for certain cancer types and a small subset of topographical features. However, previously, there has been no comprehensive evaluation that examined the effect of genome architecture and topographical features on the accumulation of somatic mutations from different mutational signatures across human cancer.

Early studies have shown that late replicating regions and condensed chromatin regions accumulate more mutations when compared to early replicating regions, actively transcribed regions, and open chromatin regions^16–19^. Subsequent analyses of hundreds of cancer genomes have revealed that differential DNA repair can explain variations in mutation rates across some cancer genomes^20^ as well as that chromatin features originating from the cell of origin, which gave rise to the tumour, can affect mutation rate and the distribution of somatic mutations^17^. Recently, Morganella *et al.* examined the effect of the genomic and the epigenomic architecture on the activity of 12 SBS signatures in breast cancer^21^. These analyses demonstrated that mutations generated by different mutational processes exhibit distinct strand asymmetries and that mutational signatures are differently affected by replication timing and nucleosome occupancy^21^. Pan-cancer exploration of strand asymmetries was also conducted for different mutation types across multiple cancer types^22^ as well as for different mutational signatures^23^. In particular, pan-cancer analyses of more than 3,000 cancers have revealed the strand asymmetries and replication timings of the 30 SBS mutational signatures from the Catalogue of Somatic Mutations in Cancer version 2 signatures database (COSMICv2)^23^. Similarly, more than 3,000 cancer genomes were used to elucidate the effect of nucleosome occupancy for the 30 substitution signatures from COSMICv2^24^. More recently, a study has also shown the interplay between the three-dimensional genome organization and the activity of certain mutational signatures^25, 26^.

Here, we report the most comprehensive evaluation of the effect of nucleosome occupancy, histone modifications, CCCTC-binding factor (CTCF) binding sites, replication timing, transcription strand asymmetry, and replication strand asymmetry on the cancer-specific accumulation of somatic mutations from distinct mutational signatures. Our analysis leverages the complete set of known COSMICv3.3 signatures (78 SBS, 11 DBS, and 18 ID) and it examines 5,120 whole-genome sequenced cancers while simultaneously utilizing 516 unique tissue-matched topographical features from the ENCODE project (**Table S1**)^27^. In all analyses, the observed patterns of somatic mutations are compared to background simulation models of mutational signatures that mimic both the trinucleotide pattern of these signatures as well as their mutational burden within each chromosome in each examined sample (**Methods**). Our results confirm many of the observations previously reported for strand asymmetry, replication timing, and nucleosome periodicity for the original COSMICv2 signatures. Further, the richer and larger COSMICv3.3 dataset allowed us to elucidate novel biological findings for some of these 30 SBS signatures revealing previously unobserved pan-cancer and cancer-specific dependencies. Additionally, this report provides the first-ever map of the genome topography of indel, doublet-base, and another 24 substitution signatures in human cancer. Moreover, our study is the first to examine the tissue-specific effect of CTCF binding and 11 different histone modifications on the accumulation of somatic mutations from different mutational signatures. As part of the results, we provide a global view of the topography of mutational signatures across 5,120 whole-genome sequenced tumours from 40 types of human cancer and we include cancer type specific examples. As part of the discussion, we zoom into two distinct case studies: *(i)* the topography of different types of clustered somatic mutations; and *(ii)* using the topography of mutational signatures to separate mutational signatures with similar patterns. Lastly, the reported results are released as part of the COSMICv3.3 signatures database, https://cancer.sanger.ac.uk/signatures, providing an unprecedented online resource for examining the topography of mutational signatures within and across human cancer types.

## RESULTS

### Transcription Strand Asymmetries

Transcription strand asymmetries have been generally attributed to transcription-coupled nucleotide excision repair (TC-NER) since bulky adducts (*e.g.*, ones due to tobacco carcinogens) in actively transcribed regions of the genome will be preferentially repaired by TC-NER^28^. Additionally, transcription-coupled damage may also lead to transcription strand asymmetry due to one of the strands being preferentially damaged during transcription^22^.

Mutational signatures with similar aetiologies generally exhibited consistent patterns of transcription strand asymmetries across cancer types. Specifically, most signatures attributed to exogenous mutational processes showed transcription strand bias with mutations usually enriched on the transcribed strand (**Figure 1*A**&E***). This included signatures SBS4/DBS2 (both previously attributed to mutagens in tobacco smoking), SBS16 (alcohol consumption), SBS24 (aflatoxin), SBS29 (tobacco chewing), SBS25/SBS31/SBS35/DBS5 (prior chemotherapy), and SBS32 (prior treatment with azathioprine). Nevertheless, for some exogenous signatures, strand asymmetries could differ between cancer types. For example, while transcriptional asymmetries for C>A and T>A mutations from SBS4 were observed across most cancer types, asymmetries for C>G mutations were only observed in lung adenocarcinoma and cancers of the head and neck (**Figure 1*C***). Interestingly, C>T mutations attributed to SBS4 had strand asymmetry only in lung adenocarcinoma. In contrast, mutational signatures due to direct damage from ultraviolet light (*viz.*, SBS7a/b/c/d and DBS1) were the only known exogenous mutational processes to exhibit transcription strand asymmetry with strong enrichment of mutations on the untranscribed strand, consistent with damage from ultraviolet light on cytosine (**Figure 1*A**&E***).

**Figure 1.**
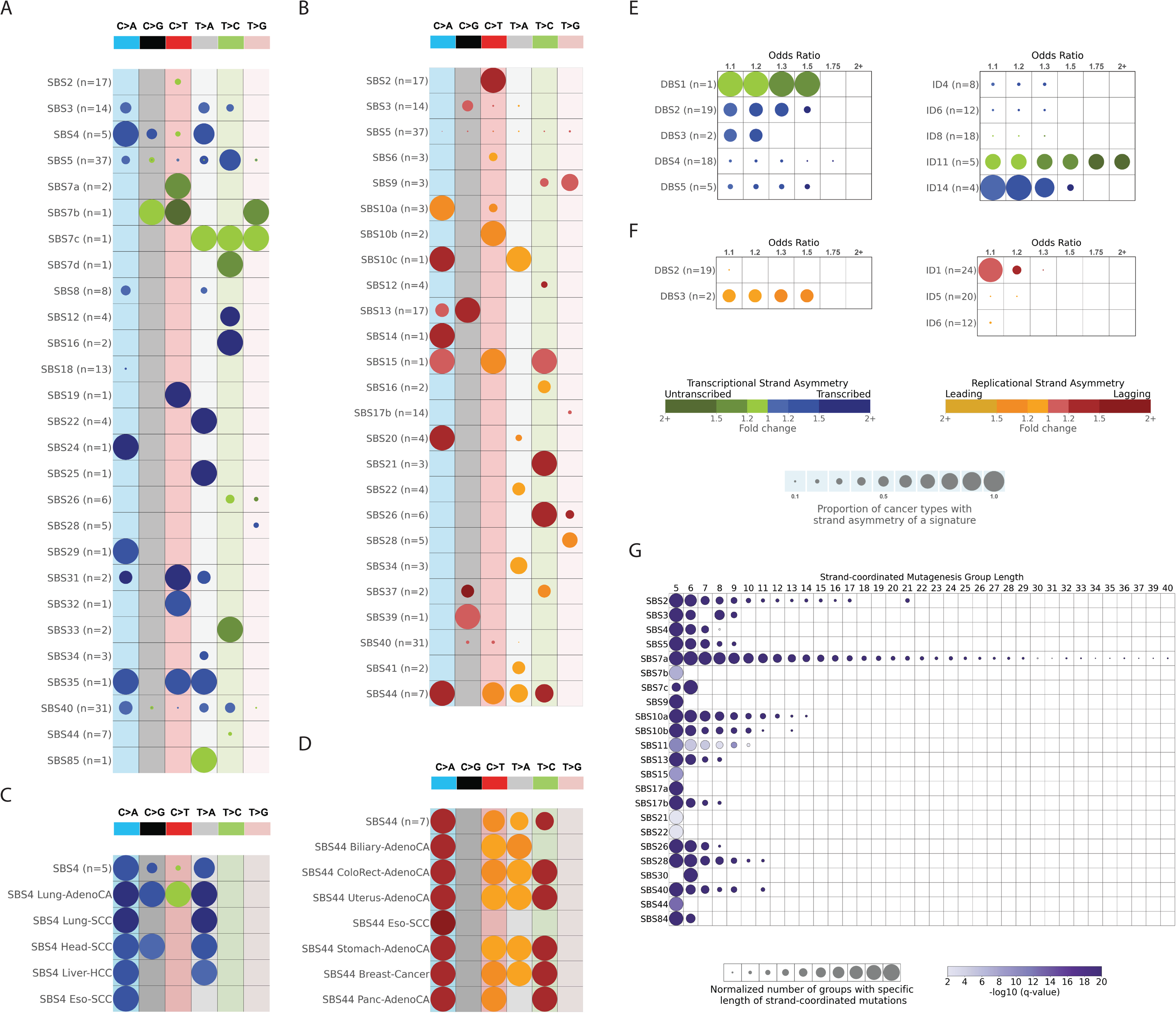
Strand asymmetries and strand-coordinated mutagenesis. ***(A)*** Transcription strand asymmetries of signatures of single base substitutions (SBSs). Rows represent the signatures, where *n* reflects the number of cancer types in which each signature was found. Columns display the six substitution subtypes based on the mutated pyrimidine base: C>A, C>G, C>T, T>A, T>C, and T>G. SBS signatures with transcription strand asymmetries on the transcribed and/or untranscribed strands with q-values ≤ 0.05 are shown in circles with blue and green colours, respectively. The colour intensity reflects the odds ratio between the ratio of real mutations and the ratio of simulated mutations, where each ratio is calculated using the number of mutations on the transcribed strand and the number of mutations on the untranscribed strand. Only odds ratios above 1.10 are shown. Circle sizes reflect the proportion of cancer types exhibiting a signature with specific transcription strand asymmetry. ***(B)*** Replication strand asymmetries of SBS signatures. Rows represent the signatures, where *n* reflects the number of cancer types in which each signature was found. Columns display the six substitution subtypes based on the mutated pyrimidine base: C>A, C>G, C>T, T>A, T>C, and T>G. SBS signatures with replicational strand asymmetries on the lagging strand or leading strand with q-values ≤ 0.05 are shown in circles with red and yellow colours, respectively. The colour intensity reflects the odds ratio between the ratio of real mutations and the ratio of simulated mutations, where each ratio is calculated using the number of mutations on the lagging strand and the number of mutations on the leading strand. Only odds ratios above 1.10 are shown. Circle sizes reflect the proportion of cancer types exhibiting a signature with specific replication strand asymmetry. ***(C)*** Transcription strand asymmetries of signatures of SBS4 across cancer types. Data are presented in a format similar to the one in panel *(A)*. ***(D)*** Replication strand asymmetries of signature SBS44 across cancer types. Data are presented in a format similar to the one in panel *(B)*. ***(E)*** Transcription strand asymmetries of signatures of doublet-base substitutions (DBSs) and of small insertions/deletions (IDs). Data are presented in a format similar to the one in panel *(A)*. ***(F)*** Replication strand asymmetries of DBS and ID mutational signatures. Data are presented in a format similar to the one in panel *(B)*. ***(G)*** Strand-coordinated mutagenesis of SBS signatures. Rows represent SBS signatures and columns reflect the lengths, in numbers of consecutive mutations, of strand-coordinated mutagenesis groups. SBS signatures with statistically significant strand-coordinated mutagenesis (q-values ≤ 0.05) are shown as circles under the respective group length with a minimum length of 5 consecutive mutations. The size of each circle reflects the number of consecutive mutation groups for the specified group length normalized for each signature. The colour of each circle reflects the statistical significance of the number of subsequent mutation groups for each group length with respect to simulated mutations.

Transcription strand asymmetry with consistent enrichment of mutations on the transcribed strand was also observed for clock-like signature SBS5 and for multiple mutational signatures with unknown aetiology, including: SBS12, SBS19, and ID14 (**Figure 1*A**&E***). Strand bias with preferences for the untranscribed strand was observed for signatures ID11 and SBS33 (both with unknown aetiology). Lastly, other mutational signatures exhibited transcription strand asymmetry in only a small subset of cancer types (**Figure 1*A**&E***).

### Mutational Signatures in Genic and Intergenic Regions

Except for SBS16 and ID11, all other signatures were enriched in intergenic regions across most cancer types with the enrichment ranging from 1.30-fold (*e.g.*, SBS24) to more than 2-fold (*e.g.*, SBS17a/b; **Figure S1*A-C***). The observed depletion of mutations in genic regions was not due to transcription strand asymmetries as correcting the asymmetries, by assigning the number of mutations on both transcribed and untranscribed strands to their highest value, resulted in only a minor alterations of the fold change increases (**Figure S1*D-E***). Overall, these results suggest that transcription strand asymmetry, usually attributed to the activity of TC-NER, does not account for the high enrichment of somatic mutations in intergenic regions.

SBS16 and ID11 showed enrichment of mutation in genic regions in liver and oesophageal cancers, while ID11 was also enriched in genic regions in cancers of the head and neck. SBS16 has been previously associated with exposure to alcohol^29–31^ and attributed to the activity of transcription-coupled damage^22^. Prior studies have also associated ID11 to alcohol consumption in oesophageal cancers^7^. Re-examining ID11 in the current cohort of whole-genome sequenced liver cancers, by comparing the mutations attributed to ID11 in 32 heavy drinkers to the mutations attributed to ID11 in 94 light drinkers, reveals a 2-fold enrichment in heavy drinkers (p-value: 1.31 x 10^-3^; Mann-Whitney U test). This and the prior associations in oesophageal cancers^7^ strongly suggest a similar exogenous mutational processes, related to alcohol consumption, accounting for the enrichment of mutation in genic regions for both signatures SBS16 and ID11.

### Replication Strand Asymmetries

Replication strand bias was consistently observed in most signatures attributed to aberrant or defective endogenous mutational processes with strand bias either on the leading or on the lagging strand (**Figure 1*B**&F***). Strong replication strand asymmetries with enrichment of mutations on the leading strand was observed for signatures previously attributed to the defective activity of polymerases, including: *(i)* SBS10a/SBS10b/DBS3 found in samples with exonuclease domain mutations in DNA polymerase epsilon (*POLE*); *(ii)* SBS9, attributed to infidelity of polymerase eta (POLH); and *(iii)* SBS10c due to defective polymerase delta (*POLD1*). Interestingly, SBS28 (unknown aetiology) exhibited a strong replication strand bias when found at high levels in POLE deficient samples.

Mutational signatures associated with defective DNA mismatch repair exhibited statistically significant replication strand bias either predominately on the leading strand (*viz.*, SBS6) or on the lagging strand (*viz.*, SBS14, SBS15, SBS20, SBS21, SBS26, SBS44, ID1). There were some minor inconsistencies of replication strand bias across cancer types. For example, SBS44 did not have replication strand asymmetry for C>T, T>A, and T>C mutations in oesophageal squamous cell carcinoma (**Figure 1*D***). Somatic mutations due to signatures SBS2 and SBS13, both attributed to the aberrant behaviour of the APOBEC3 family of deaminases^32^, were found enriched on the lagging strand in all cancer types. This result is consistent with the observation that single-stranded DNA formed during DNA replication on the lagging strand is a major substrate for the APOBEC3 family of deaminases^33, 34^. Lastly, several other mutational signatures, most with unknown aetiology, exhibited replication strand bias within a small set of cancer types (**Figure 1*B**&F***).

### Strand-coordinated Mutagenesis

Prior analyses have shown that certain types of mutations on the same reference allele were observed on the same strand more frequently than expected by chance^21, 34, 35^. This strand-coordinated clustered mutations usually arise due to damage on single-stranded DNA, and they are often indicative of the formation of hypermutable loci in the genome^33, 34^.

SBS7a, attributed to ultraviolet (UV) light, attained the highest strand-coordinated mutagenesis with lengths of subsequent mutations up to 40 consecutive mutations (**Figure 1*G***). In contrast, other mutational signatures attributed to ultraviolet light, mainly, SBS7b/c/d, either did not exhibit or exhibited much lower strand-coordinated mutagenesis. APOBEC3-attributed SBS2 and SBS13 showed strand-coordinated mutagenesis with as many as 21 consecutive strand-coordinated mutations. Additionally, strand-coordinated mutations were observed for SBS17b (unknown aetiology), SBS10a/b (*POLE* deficiency), SBS4 (tobacco smoking), SBS26 (defective mismatch repair), and SBS28 (unknown aetiology).

### The Effect of DNA Replication Timing

Consistent with prior reports^18, 36–38^, the aggregated set of somatic mutations was shown to be enriched in late replicating regions for most cancer types (**Figure 2*A***). Specifically, from the examined 40 cancer types, SBSs were found more common in regions of the genome that undergo late replication in 39/40 cancer types and were not associated with replication only in uveal melanoma (**Figure 2*A***). Similarly, DBSs and IDs were enriched in late replicating regions in 18/18 and 30/32 cancer types, respectively. Note that due to their lower mutational burdens, we could confidently evaluate DBSs and IDs only in a subset of cancer types. In agreement with the aggregates analysis, most mutational signatures imprinted somatic mutations with an increased normalized mutational density from early to late replicating regions (**Figure S2**). For example, SBS3 (defective homologous recombination) was enriched in late replicating regions in all 14 cancer types where the signature can be confidently evaluated. Other examples include signatures DBS2 and ID1, which were also consistently enriched in all examined cancer types (**Figure 2*B***).

**Figure 2.**
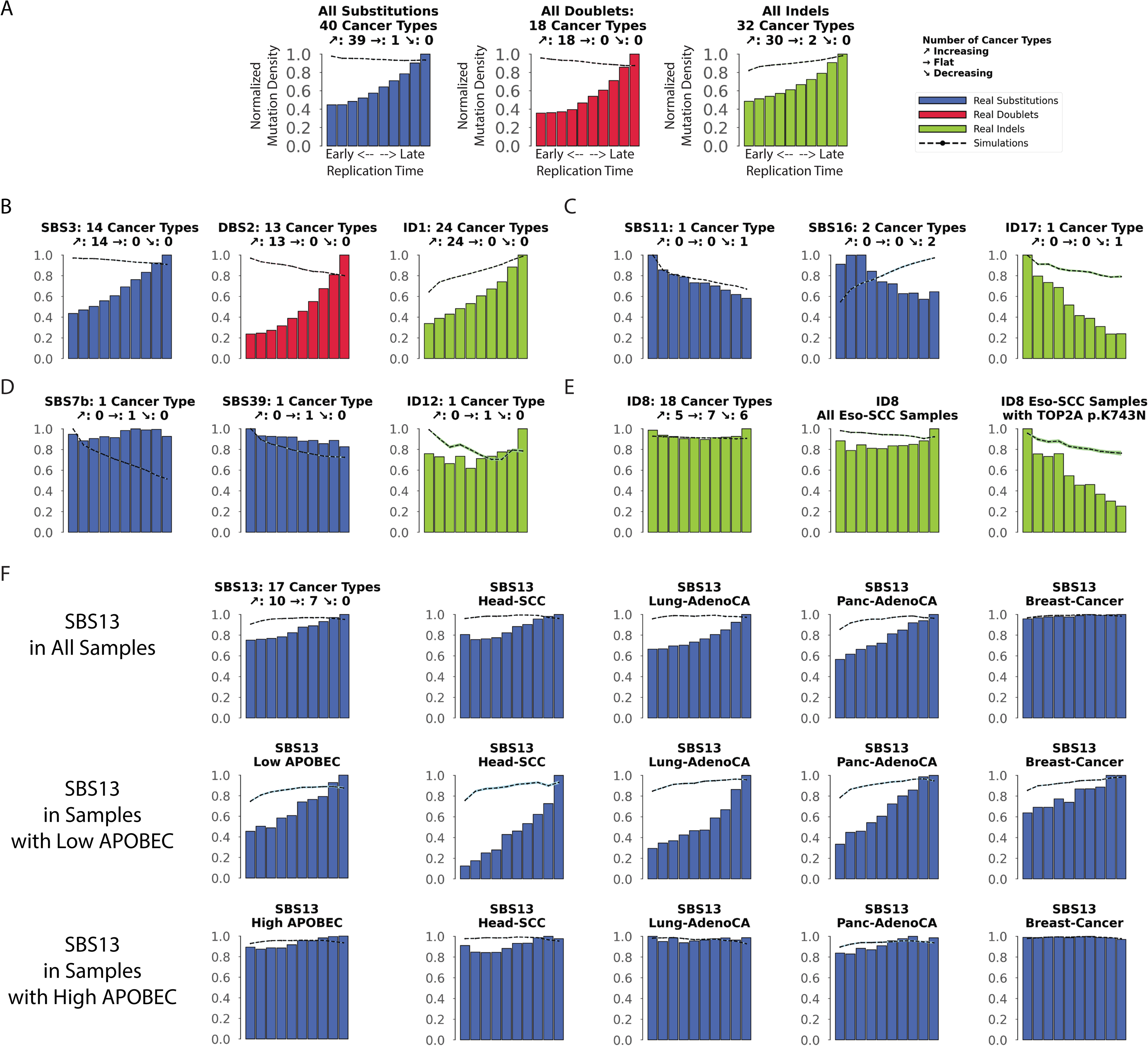
Interplay between replication timing and mutational signatures. Replication time data are separated into deciles, with each segment containing exactly 10% of the observed replication time signal (x-axes). Normalized mutation densities per decile (y-axes) are presented for early (left) to late (right) replication domains. Real data for single base substitutions (SBSs) signatures are shown as blue bars, for doublet-base substitutions (DBSs) signatures as red bars, and for small insertions/deletions (IDs) signatures as green bars. Simulated somatic mutations are shown as dashed lines. Where applicable, the total number of evaluated cancer types for a particular mutational signature is shown on top of each plot (*e.g.,* 18 cancer types were evaluated for ID8 in panel E). For each signature, the number of cancer types where the mutation density increases with replication timing is shown next to ⬈ (*e.g.,* 5 cancer types for ID8). Similarly, the number of cancer types where the mutation density decreases with replication timing is shown next to ⬊ (*e.g.,* 6 cancer types for ID8). Lastly, the number of cancer types where the mutation density is not affected by replication timing is shown next to ➞ (*e.g.,* 7 cancer types for ID8). ***(A)*** All SBSs, DBSs, and IDs across all examined cancer types with each cancer type examined separately. ***(B)*** Exemplar signatures consistently associated with late replication timing. ***(C)*** Exemplar signatures consistently associated with early replication timing. ***(D)*** Exemplar signatures consistently unaffected by replication timing. ***(E)*** ID8 as a mutational signature inconsistently affected by replication timing. ***(F)*** The effect of replication timing on APOBEC3-associated signature SBS13 in samples with low and high APOBEC3 mutational burden.

Nevertheless, at least seven mutational signatures were found predominately enriched in early replicating regions, including: ID17 likely due to *TOP2A* mutations; SBS11 due to temozolomide therapy; SBS16, ID11 both associated with alcohol consumption; SBS6, SBS15 both attributed to mismatch repair deficiency; and SBS84 due to the aberrant activities of activation-induced (AID) cytidine deaminases (**Figure 2*C***; **Figure S2**). Moreover, multiple mutational signatures were generally unaffected by replication timing, including: SBS7b (UV-light); SBS20, SBS21, SBS44 (attributed to failure of mismatch repair); SBS30 (deficient base excision repair); and SBS39, ID12 (unknown aetiology; **Figure 2*D***; **Figure S2**). The lack of association with replication timing for some of these mutational signatures can be potentially attributed to the activity of DNA translesion polymerases^39, 40^.

Interestingly, a number of mutational signatures exhibited cancer-type specific associations with replication timing (**Figure S2**). For example, signature ID8 was enriched with replication timing in 5 cancer types, depleted in 6 cancer types, and unaffected by replication timing in 7 cancer types (**Figure 2*E***). Multiple aetiologies have been associated with ID8^4, 41^, including mutations resulting in K743N amino acid change in TOP2A. All samples harbouring such mutations in *TOP2A* exhibited an enrichment of ID8 in early replicating regions (**Figure 2*E***). The other cancer-type specific mechanisms resulting in different associations with replication timing for ID8 remain unknown.

Another notable example of cancer-type specific associations with replication timing is the APOBEC3-associated SBS13 (**Figure S2**). SBS13 showed no dependence with replication timing in 7/17 cancer types (*viz.*, bladder, breast, uterus, cervix, ovary, thyroid, and acute lymphocytic leukaemia; **Figure 2*F***). This behaviour is consistent with prior reports where SBS13 was attributed to uracil excision of deaminated cytosine followed by processing by DNA translesion polymerases in breast cancer^39, 40^. Surprisingly, in 10/17 cancer types, SBS13 was highly enriched in late replicating regions. Using a previously defined approach for separating the cancer samples on ones where SBS13 is not a hypermutator (low APOBEC3) and ones where SBS13 is a hypermutator (high APOBEC3) revealed that the lack of dependence with replication timing is predominately characteristic for hypermutated samples (**Figure 2*F***). This result indicates that DNA translesion polymerases may play a significantly larger role in APOBEC3 hypermutators than previously anticipated.

### The Effect of Nucleosome Occupancy

Nucleosomes are the basic packing units of chromatin with each nucleosome consisting of ∼147 base-pair (bp) DNA wrapped around a histone octamer with 60 to 80 bp linker DNA between consecutive nucleosomes^42, 43^. Previous analyses have revealed dependencies between mutational signatures operative in breast cancer and nucleosome occupancy^21^ as well as a pan-cancer periodicity of mutation rates within nucleosomes due to multiple substitution signatures^24^.

However, beyond breast cancer, there has been no cancer-specific examination of the effect of nucleosome occupancy on different mutational signatures.

Aggregated somatic mutations and mutations attributed to most mutational signatures were depleted near nucleosomes compared to simulated data that mimic the mutational landscapes of the examined cancer genomes (**Figure 3*A***). Remarkably, the majority of SBS, DBS, and ID mutational signatures were similarly affected by nucleosome occupancy across most cancer types (**Figure S3**). Some signatures were consistently enriched in the vicinity of nucleosomes. For example, clock-like signature SBS1 exhibited a pattern closely mimicking simulated data and showing higher number of mutations at nucleosomes in 36/36 cancer types, including cancers of the lung, head and neck, liver, and oesophagus (**Figure 3*B***). In contrast, some signatures were markedly different that simulated data (**Figure S3**), including signature DBS2 which was consistently depleted across 13/13 cancer types (**Figure 3*C***). Moreover, some signatures were depleted in nucleosomes and strikingly appeared at linker DNA (**Figure S3**). For example, clock-like signature ID1 was depleted when compared to simulated data and it exhibited depletion in nucleosomes in 24/24 examined cancer types (**Figure 3*D***). The mutations engraved by most flat mutational signatures (*e.g.*, SBS5, SBS8, SBS40) were generally unaffected by nucleosomes (**Figure S3**).

**Figure 3.**
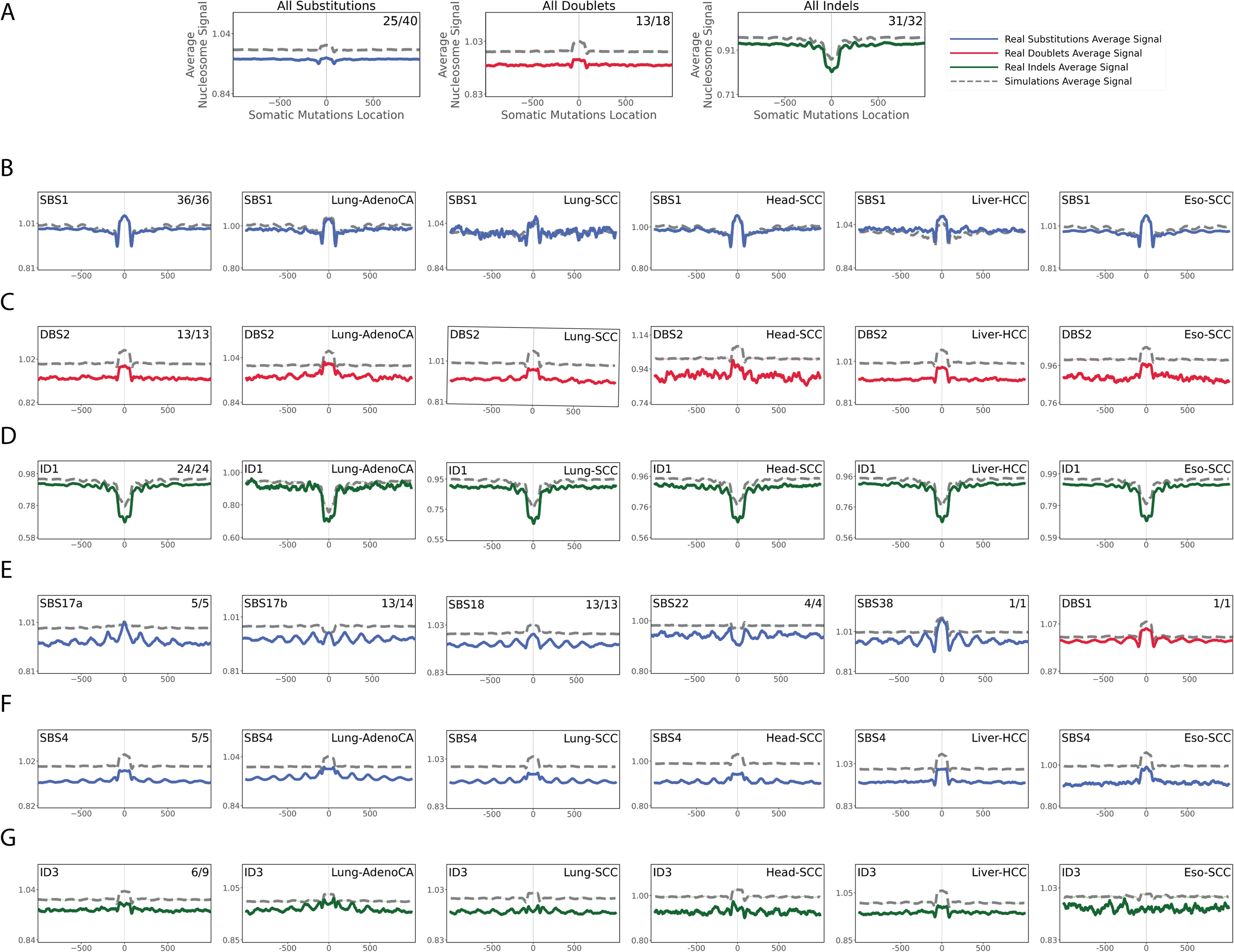
Relationship between mutational signatures and nucleosome occupancy. In all cases, solid lines correspond to real somatic mutations with blue solid lines reflecting single base substitutions (SBSs), red doublet-base substitutions (DBSs), and green small insertions/deletions (IDs). Simulated somatic mutations are shown as dashed lines. Solid lines and dashed lines display the average nucleosome signal (y-axes) along a 2 kilobase window (x-axes) centred at the mutation start site for real and simulated mutations, respectively. The mutation site is annotated in the middle of each plot and denoted as 0. The 2 kilobase window encompasses 1,000 base-pairs 5’ adjacent to each mutation as well as 1,000 base-pairs 3’ adjacent to each mutation. Where applicable, the total number of similar and considered cancer types using an *X/Y* format, with *X* being the number of cancer types where a signature has similar nucleosome behaviour (Pearson correlation ≥ 0.5 and q-value ≤ 0.05) and *Y* representing the total number of examined cancer types for that signature. For example, signature ID3 in panel G annotated with 6/9 reflects a total of 9 examined cancer types with similar nucleosome behaviour observed in 6 cancer types. ***(A)*** All SBSs, DBSs, and IDs across all examined cancer types with each cancer type examined separately. The nucleosome occupancy of signatures SBS1 ***(B)***, DBS2 ***(C)***, and ID1 ***(D)*** are shown across all cancer types as well as within cancers of the lung, head and neck, liver, and oesophagus. ***(E)*** Signatures with consistent periodicities of mutation rates around the nucleosome. Tobacco-associated SBS4 ***(F)*** and ID3 ***(G)*** exhibiting periodicities of mutation rates only in certain cancer types.

Different types of periodicities of mutation rates around the nucleosome structure were observed for signatures associated with tobacco smoking (SBS4 and ID3), ultraviolet light (SBS7a/b/c/d), POLE deficiency (SBS10a), aristolochic acid (SBS22), and reactive oxygen species (SBS18, SBS36, SBS38; **Figures 3*E* & S3**). Interestingly, signatures SBS17a/b also showed similar periodic dependencies (**Figure 3*E*)** providing further circumstantial evidence for the hypothesis that SBS17a/b may also be due to reactive oxygen species damage of the deoxyribonucleoside triphosphate pools^23, 44–48^. With the exception of signature SBS22 and ID3, all other periodic signatures exhibited enrichment of mutations at nucleosomes (**Figure 3*E**&G***). Further, for most signatures periodicity of mutation rates was observed in each cancer types where the signature was operative (**Figure S3)**. Nevertheless, signature SBS4 showed strong periodicity in cancers of the lung and head and neck but not in cancers of the liver or oesophagus (**Figure 3*F***). Similarly, signature ID3 exhibited periodic behaviour only in cancers of the lung but not in any other cancer type (**Figure 3*G***).

### The Effect of CTCF Binding

CCCTC-binding factor (CTCF) is a multi-purpose sequence-specific DNA-binding protein with an essential role in transcriptional regulation, somatic recombination, and chromatin architecture^49^. The human genome harbours many CTCF binding sites with prior studies reporting that mutations due to ultraviolet light are enriched in CTCF binding sites^50^.

Somatic mutations exhibited clear patterns of both enrichment and/or periodicity for multiple mutational signatures and CTCF binding sites (**Figure 4**). While some signatures were consistently depleted at CTCF biding sites across the majority of cancer types when compared to simulated data (SBS1, SBS9, SBS10a/b, SBS15, SBS37, SBS84, and SBS85), others were commonly enriched (SBS3, SBS5, SBS7a/b/d, SBS12, SBS17a/b, SBS18, SBS22, and SBS40; DBS1; ID5, ID6, ID8, and ID9; **Figure 4*A***).

**Figure 4.**
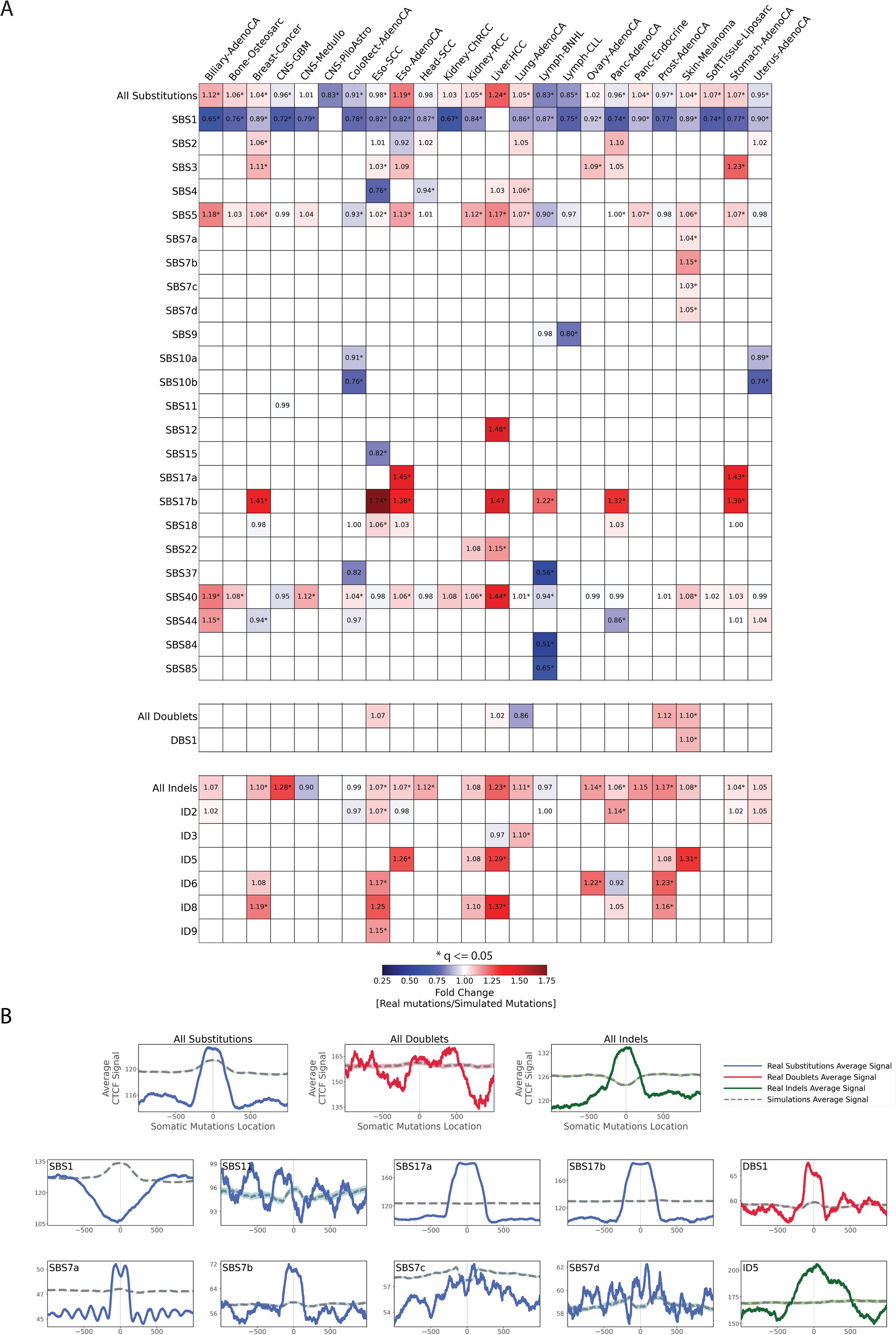
Relationship between mutational signatures and CTCF binding sites. ***(A)*** Enrichments and depletions of somatic mutations within CTCF binding sites. Heatmaps display only mutational signatures and cancer types that have at least one statistically significant enrichment or depletion of somatic mutations attributed to signatures of either single base substitutions (SBSs), doublet-base substitutions (DBSs), or small insertions/deletions (IDs). Red colours correspond to enrichments of real somatic mutations when compared to simulated data. Blue colours correspond to depletions of real somatic mutations when compared to simulated data. The intensities of red and blue colours reflect the degree of enrichments or depletions based on the fold change. White colours correspond to lack of data for performing statistical comparisons *(e.g.*, signature not being detected in a cancer type). Statistically significant enrichments and depletions are annotated with * (q-value ≤ 0.05). ***(B)*** The top three panels reflect average CTCF occupancy signal for all SBSs, DBS, and IDs across all examined cancer types. Bottom panels reflect all somatic mutations attributed for several exemplar mutational signatures across all cancer types. In all cases, solid lines correspond to real somatic mutations with blue solid lines reflecting SBSs, red solid lines reflecting DBSs, and green solid lines reflecting IDs. Solid lines and dashed lines display the average CTCF binding signal (y-axes) along a 2 kilobase window (x-axes) centred at the mutation start site for real and simulated mutations, respectively. The mutation start site is annotated in the middle of each plot and denoted as 0. The 2 kilobase window encompasses 1,000 base-pairs 5’ adjacent to each mutation as well as 1,000 base-pairs 3’ adjacent to each mutation.

Aggregated single base substitutions exhibited an inconsistent behaviour across cancer types with enrichment in some cancers (*e.g.,* liver cancers) and depletions in others (*e.g.*, lymphomas). In contrast, indels were enriched at CTCF binding sites in the majority of cancer types (**Figure 4*A***). Remarkably, the effect of CTCF occupancy tended to be also consistent for many signatures with similar aetiologies. Strong periodicities of mutation rates around CTCF binding sites were observed for UV-associated signature SBS7a but not for UV-associated signatures DBS1 and SBS7b/c/d (**Figure 4*B***).

Mutations due to SBS9, associated with defective polymerase eta driven replication errors, and signatures SBS10a/b, found in samples with mutations in *POLE* and/or *POLD1*, were strikingly depleted at CTCF binding sites. Signatures SBS15, associated microsatellite instability, was strongly depleted at CTCF binding sites (**Figure 4*A***).

Only one of the clock-like signatures, SBS1, exhibited a depletion of mutations at CTCF binding sites (**Figure 4*A***) while simulated data indicated that SBS1 should be enriched at these sites (**Figure 4*B***). Signature SBS3, attributed to defective homologous recombination, was highly elevated in CTCF binding sites for breast, ovarian, stomach, and oesophageal cancers. Signatures SBS17a/b exhibited a striking enrichment at CTCF binding sites in all cancer types with sufficient number of mutations from each signature (**Figure 4*A***). SBS17a showed enrichment in stomach and oesophageal cancers, while SBS17b shows enrichment for stomach, oesophageal, breast, pancreatic cancers, and non-Hodgkin’s lymphomas. In contrast, simulated data indicate that CTCF binding should have no effect on the accumulation of mutations from signatures SBS17a/b (**Figure 4*B***).

### The Effect of Histone Modifications

Each nucleosome consists of four pairs of core histones: H2A, H2B, H3, and H4. Post-translational modifications of histone tails play a key role in regulating DNA replication, gene transcription, and DNA damage response^51^. To evaluate the effect of histone modifications on the accumulation of mutations from different mutational signatures, we mapped the depletion or enrichment of mutations compared to simulated data in the context of the tissue specific positions of 11 histone modifications: *(i)* H2AFZ, a replication-independent member of the histone H2A family that renders chromatin accessible at enhancers and promoters regulating transcriptional activation and repression^52^; *(ii)* H3K4me1, histone mark often associated with enhancer activity^53^; *(iii)* H3K4me2, a histone post-translational modification enriched in *cis*-regulatory regions, including both enhancers and promoters^54^; *(iv)* H3K4me3, post-translational modification enriched in active promoters near transcription start sites^55^; *(v)* H3K9ac, associated with active gene promoters and active transcription^56^; *(vi)* H3K9me3, silencer, typical mark of constitutive heterochromatin^57^; *(vii)* H3K27ac, histone modification generally contained at nucleosomes flanking enhancers^55^; *(viii)* H3K27me3, repressive, associated with silent genes^58^; *(ix)* H3K36me3, associated with transcribed regions and playing a role in regulating DNA damage repair^59^; *(x)* H3K79me2, detected in the transcribed regions of active genes^60^; and *(xi)* H4K20me1, found in gene promoters and associated with gene transcriptional elongation and transcription activation^61^.

Aggregated substitutions, dinucleotides, and indels exhibited dissimilar behaviour for different histone modifications across cancer types (**Figure S4**). Aggregated substitutions were predominately depleted around H2AFZ, H3K4me2, H3K4me3, and H3K27ac in approximately half of the examined cancer types (**Figure S4*A-C***). Aggregated doublets and indels did not have any clear pan-cancer preference but showed cancer-type specific enrichments and depletions. In contrast, majority of mutational signatures had generally similar behaviour in vicinity of different histone modifications revealing that histone modifications have similar effect on mutagenesis across cancer types (**Figure S4**). Most SBS mutational signatures were either unaffected or depleted near histone marks (**Figure S4*A***). Notable exceptions were APOBEC3-associated signatures SBS2 and SBS13, AID-associated signatures SBS84 and SBS85, and *POLH* attributed SBS9 which were generally enriched near most histone modifications (**Figure S4*A***). Doublet signatures DBS1, DBS2, DBS3, DBS4, and DBS5 were also predominately depleted near most histone marks (**Figure S4*B***). In contrast, signatures DBS7, DBS9, and DBS11 were highly enriched near most histone marks. Most ID mutational signatures were either unaffected or very highly enriched near histone marks (**Figure S4*C***) with the only exceptions being depletions of: *(i)* ID1 and ID6 near H2AZ; *(ii)* ID3 in the vicinity of H3K4me3; *(iii)* ID5 near H3K27me3; and *(iv)* ID14 in the vicinity H3K36me3. While enrichments and depletions of somatic mutations in the vicinity of histone marks were commonly observed for different mutational signatures (**Figure S4*A-C***), there was no specific pattern of mutations within 1,000 base-pairs for any of the examined histone modifications. Exemplars of typically observed patterns of enrichments, depletions, or no changes around different histone modifications are provided for signatures SBS7a and ID1 across several histone modifications (**Figure S5*D***).

Next, we examine two mutational signatures that exhibited inconsistent enrichments and depletions near specific histone marks. Clock-like signature SBS1 was consistently depleted across cancer-types for multiple histone marks, including H3K9me3 (**Figure 5*A***). Nevertheless, SBS1 exhibited enrichment of mutations near H3K9me3 in two cancer types of the central nervous system, depletion of mutations near H3K9me3 in three haematological malignancies, and no effect in all other solid tumour types (**Figure 5*A***). Similarly, signature ID1 exhibited dissimilar behaviour near H3K27ac with enrichments in medulloblastoma and lymphoma, depletions in stomach and prostate cancer, and no change in most other cancer types (**Figure 5*B***).

**Figure 5.**
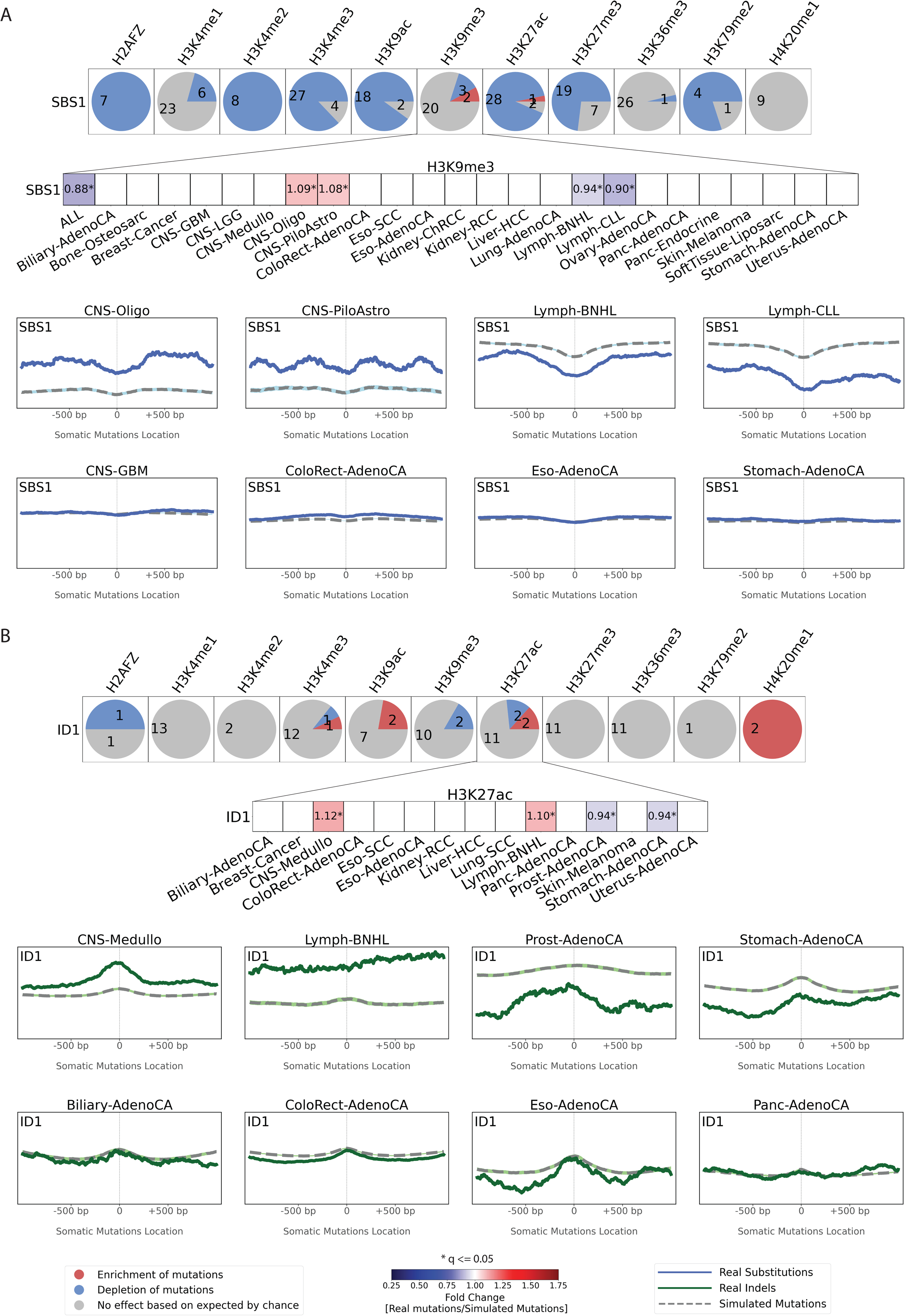
Exemplar relationships between mutational signatures and histone modifications. The effect of histone modifications is shown for signatures SBS1 ***(A)*** and ID1 ***(B)***. For each signature, an evaluation was made for each of the 11 histone marks across all cancer types with sufficient numbers of somatic mutations with results presented as circles. Each circle is separated in red, blue, and grey segments proportionate to the cancer types in which the signature has a specific behaviour. A red segment in a circle reflects the signature being enriched in the vicinity of a histone modification (q-value ≤ 0.05 and at least 5% enrichment). A blue segment in a circle reflects the signature being depleted in the vicinity of a histone modification (q-value ≤ 0.05 and at least 5% depletion). A grey segment in a circle corresponds to neither depletion nor enrichment of the signature in the vicinity of a histone modification. The figure zooms into the effect of H3K9me3 on SBS1 ***(A)*** and H3K27ac on ID1 ***(B)***. Red colours correspond to enrichments of real somatic mutations when compared to simulated data. Blue colours correspond to depletions of real somatic mutations when compared to simulated data. The intensities of red and blue colours reflect the degree of enrichments or depletions based on the fold change. We further zoom into the patterns of mutations around H3K9me3 and H3K27ac. Solid lines correspond to real somatic mutations with blue solid lines reflecting SBSs and green solid lines reflecting IDs. Solid lines and dashed lines display the average histone mark signal (y-axes) along a 2 kilobase window (x-axes) centred at the mutation start site for real and simulated mutations, respectively. The mutation start site is annotated in the middle of each plot and denoted as 0. The 2 kilobase window encompasses 1,000 base-pairs 5’ adjacent to each mutation as well as 1,000 base-pairs 3’ adjacent to each mutation.

## DISCUSSION

Our analysis provides a comprehensive resource that maps the effects of topographical genomic features on the cancer-specific accumulation of somatic mutations from distinct mutational signatures. The reported results confirmed many of the prior observations for strand asymmetry, replication timing, and nucleosome periodicity for some of the original 30 COSMICv2 SBS signatures^21, 23, 24^. The examined larger dataset provided us with a greater resolution to identify previously unobserved pan-cancer and cancer-specific dependencies for some of these 30 signatures as well as to reveal the effect of genome architecture on the accumulation of another 46 mutational signatures across human cancer. Importantly, this report also provides the first-ever examination of the tissue-specific effect of CTCF binding and 11 different histone modifications on the accumulation of somatic mutations from different mutational signatures. In addition to the comprehensive global view in the results section, in this discussion, we zoom into two specific case studies to further illustrate the power of using this resource for examining topography of mutational signatures.

First, analysis of SBS28 in *POLE* deficient samples (*POLE^-^*) and *POLE* proficient samples (*POLE^+^*) revealed a distinct behaviour (**Figure 6**). While the trinucleotide patterns of SBS28 in *POLE^+^*and *POLE^-^* samples were similar (cosine similarity: 0.96), SBS28 in POLE^-^ samples accounted for 97.7% mutations of all SBS28 mutations and it exhibited a clear enrichment in late replicating regions as well as depletions at nucleosomes and at CTCF binding sites (**Figure 6*B**-D*,*F***). Moreover, SBS28 in *POLE^-^* samples showed a strong replication strand bias on the leading strand and exhibited a strand-coordinated mutagenesis with as many as 11 consecutively mutated substitutions (**Figure 6*E*,*G***). In contrast, SBS28 in *POLE^+^* samples were enriched in early replication regions, lacked depletion of mutations at nucleosomes or CTCF binding sites, had weak replication strand bias on lagging strand, and did not exhibit much of a strand-coordinated mutagenesis (**Figure 6**). Based on these topographical differences, we have now split SBS28 into two distinct signatures: *(i)* SBS28a due to *POLE* deficiency found in ultra-hypermutate colorectal and uterine cancers; and *(ii)* SBS28b with unknown aetiology found in lung and stomach cancers.

**Figure 6.**
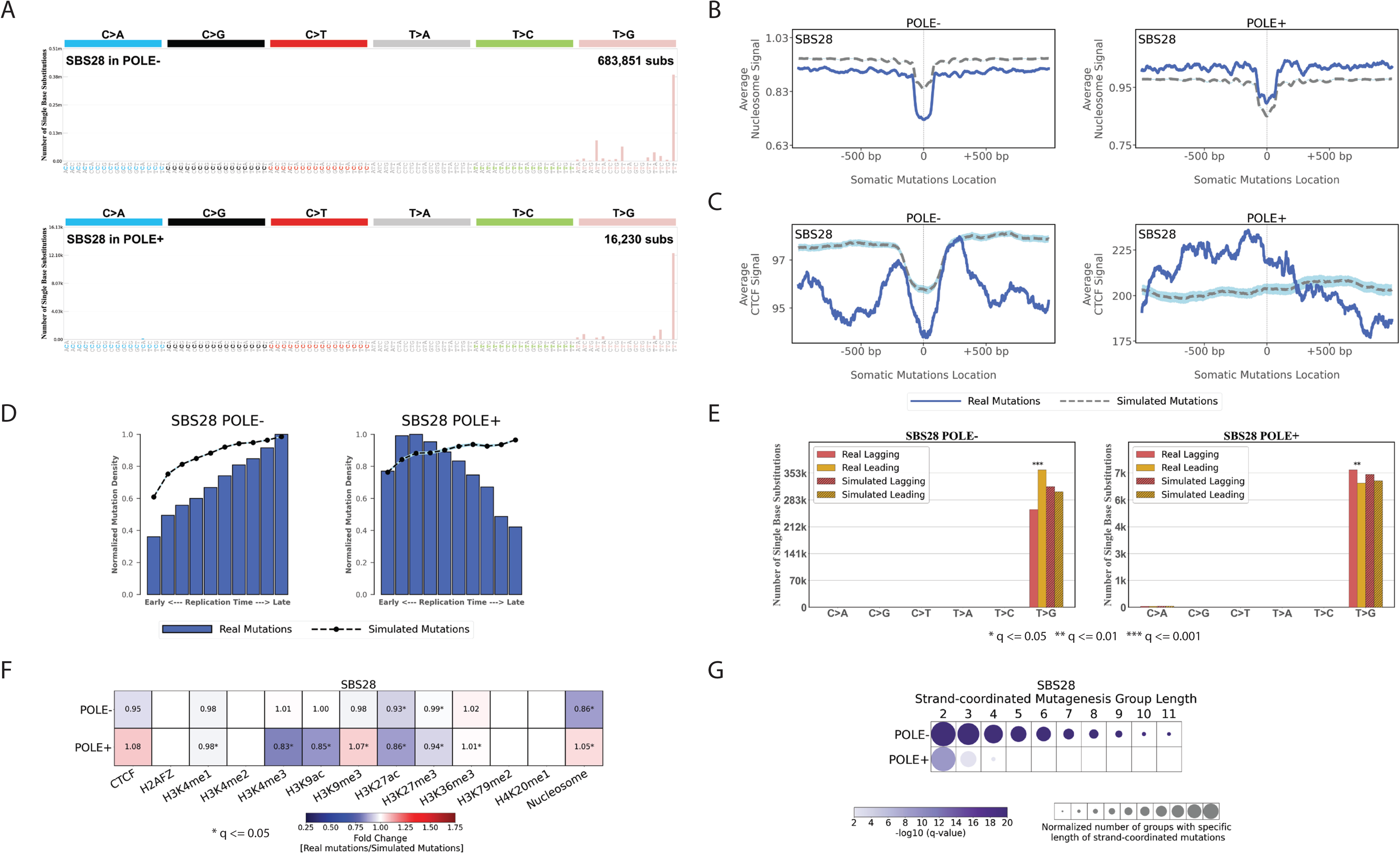
Topography of signature SBS28 in *POLE* deficient (*POLE^-^*) and *POLE* proficient (*POLE^+^*) samples. ***(A)*** Mutational patterns of signature SBS28 in *POLE*^-^ and *POLE^+^* samples displayed using the conventional 96 mutational classification schema for single base substitutions. ***(B)*** Nucleosome occupancy of SBS28 in *POLE^-^* and *POLE^+^* samples. Blue solid lines and grey dashed lines display the average nucleosome signal (y-axes) along a 2 kilobase window (x-axes) centred at the mutation start site for real and simulated mutations, respectively. The mutation start site is annotated in the middle of each plot and denoted as 0. The 2 kilobase window encompasses 1,000 base-pairs 5’ adjacent to each mutation as well as 1,000 base-pairs 3’ adjacent to each mutation. ***(C)*** CTCF occupancy of SBS28 in *POLE^-^* and *POLE^+^* samples. Blue solid lines and grey dashed lines display the average CTCF binding signal (y-axes) along a 2 kilobase window (x-axes) centred at the mutation start site for real and simulated mutations, respectively. The mutation start site is annotated in the middle of each plot and denoted as 0. The 2 kilobase window encompasses 1,000 base-pairs 5’ adjacent to each mutation as well as 1,000 base-pairs 3’ adjacent to each mutation. ***(D)*** Replication timing of SBS28 mutations in *POLE^-^* and *POLE^+^* samples. Replication time data are separated into deciles, with each segment containing exactly 10% of the observed replication time signal (x-axes). Normalized mutation densities per decile (y-axes) are presented for early (left) to late (right) replication domains. Normalized mutation densities of real somatic mutations and simulated somatic mutations from early to late replicating regions are shown as blue bars and dashed lines, respectively. ***(E)*** Replication strand asymmetry of SBS28 mutations in *POLE^-^*and *POLE^+^* samples. Bar plots display the number of mutations accumulated on the lagging strand and leading strand for six substitution subtypes based on the mutated pyrimidine base: C>A, C>G, C>T, T>A, T>C, and T>G in red and yellow colours, respectively. Simulated mutations on lagging and leading strands are displayed in shaded bar plots. Statistically significant strand asymmetries are shown with stars: * q-value ≤ 0.05; ** q-value ≤ 0.01; *** q-value ≤ 0.001. ***(F)*** Enrichments and depletions of SBS28 somatic mutations in *POLE^-^* and *POLE^+^* samples within CTCF binding sites, histone modifications, and nucleosome occupied regions. Red colours correspond to enrichments of real somatic mutations when compared to simulated data. Blue colours correspond to depletions of real somatic mutations when compared to simulated data. The intensities of red and blue colours reflect the degree of enrichments or depletions based on the fold change. White colours correspond to lack of data for performing statistical comparisons. Statistically significant enrichments and depletions are annotated with * (q-value ≤ 0.05). ***(G)*** Strand-coordinated mutagenesis of SBS28 mutations in *POLE^-^*and *POLE^+^* samples. Rows represent SBS28 signature in *POLE^-^*and *POLE^+^* samples across all cancer types and columns reflect the lengths, in numbers of consecutive mutations, of strand-coordinated mutagenesis groups. Statistically significant strand-coordinated mutagenesis (q-value ≤ 0.05) are shown as circles under the respective group length with a length starting from 2 to 11 consecutive mutations. The size of each circle reflects the number of consecutive mutation groups for the specified group length normalized for each SBS28 signature in *POLE^-^* and *POLE^+^* samples. The colour of each circle reflects the statistical significance of the number of subsequent mutation groups for each group length with respect to the simulated mutations using –log_10_ (q-value).

Second, our analyses revealed striking difference in topographical features of clustered and non-clustered somatic mutations in 288 whole-genome sequenced B-cell malignancies^4^. In particular, the topographical behaviours of single base substitutions were examined after separating them into non-clustered mutations, diffuse hypermutation of substitutions termed *omikli*^62^, and longer clusters of strand-coordinated substitutions termed *kataegis*^34, 35, 63^. In contrast to most cancer types, where *omikli* and *kataegis* are predominately generated by APOBEC3 deaminases^64^, in B-cell malignancies, these clustered events are almost exclusively imprinted by the activity of AID^64^. Further, the overall pattern of non-clustered mutations was very different than the ones of *omikli* or *kataegis*. A representative example is provided using a single malignant B-cell lymphoma (**Figure 7*A***) where non-clustered and clustered mutations have very different trinucleotide patterns (**Figure 7*B*–*D***). Non-clustered mutations exhibited different topographical features when compared to *omikli* or *kataegis*. Specifically, while non-clustered mutations had some minor periodicity in regard to nucleosome occupancy, such periodicity was not observed for any type of clustered events (**Figure 7*E***). Similarly, non-clustered mutations were slightly depleted around CTCF binding sites while *omikli* and *kataegis* were very highly depleted (**Figure 7*F**&H***). Further, non-clustered and *omikli* events were clearly enriched in late replication regions while *kataegis* was highly enriched in early replication regions (**Figure 7*G***). Distinct patterns of enrichments were also observed for both *omikli* and *kataegis* mutations in the vicinity of promoter and enhancer sites delineated by histone marks of H3K4me3, H3K9ac, H3K27ac, H3K36me3, and H4K20me1(**Figure 7*H***). Only very minor differences were observed for transcription or replication strand asymmetries between clustered and non-clustered somatic mutations across the 288 whole-genome sequenced B-cell malignancies (**Figure S5**).

**Figure 7.**
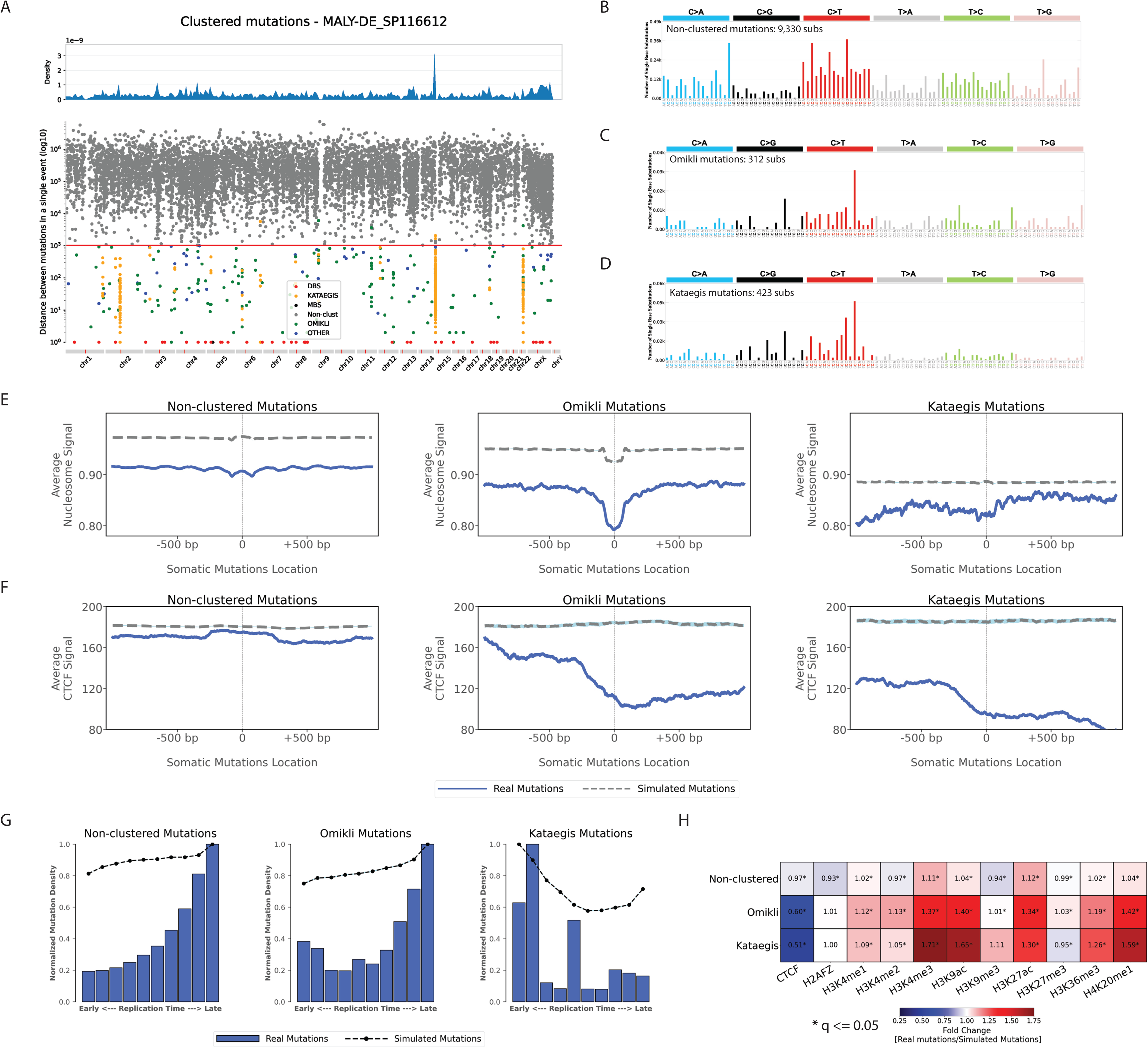
Topography of non-clustered, *omikli*, and *kataegis* substitutions across 288 whole-genome sequenced B-cell malignancies. ***(A)*** A rainfall plot of an example B-cell malignancy sample, MALY-DE_SP116612, depicting the intra-mutational distance (IMD) distributions of substitutions across genomic coordinates. Each dot represents the minimum distance between two adjacent mutations. Dots are coloured based on their corresponding classifications. Specifically, non-clustered mutations are shown in grey, doublet-base substitutions (DBSs) in red, multi-base substitutions (MBSs) in black, *omikli* events in green, *kataegis* events in orange, and all other clustered events in blue. The red line depicts the sample-dependent IMD threshold for each sample. Specific clustered mutations may be above this threshold due to corrections for regional mutation density. ***(B-D)*** The trinucleotide mutational spectra for the different catalogues of non-clustered, *omikli,* and *kataegis* mutations for the exemplar sample (DBSs and MBSs are not shown). ***(E)*** Nucleosome occupancy of non-clustered, *omikli*, and *kataegis* mutations of B-cell malignancies. Blue solid lines and grey dashed lines display the average nucleosome signal (y-axes) along a 2 kilobase window (x-axes) centred at the mutation start site for real and simulated mutations, respectively. The mutation start site is annotated in the middle of each plot and denoted as 0. The 2 kilobase window encompasses 1,000 base-pairs 5’ adjacent to each mutation as well as 1,000 base-pairs 3’ adjacent to each mutation. ***(F)*** CTCF occupancy of non-clustered, *omikli*, and *kataegis* mutations of B-cell malignancies. Blue solid lines and grey dashed lines display the average CTCF signal (y-axes) along a 2 kilobase window (x-axes) centred at the mutation start site for real and simulated mutations, respectively. The mutation start site is annotated in the middle of each plot and denoted as 0. The 2 kilobase window encompasses 1,000 base-pairs 5’ adjacent to each mutation as well as 1,000 base-pairs 3’ adjacent to each mutation. ***(G)*** Replication timing of non-clustered, *omikli*, and *kataegis* mutations of B-cell malignancies. Replication time data are separated into deciles, with each segment containing exactly 10% of the observed replication time signal (x-axes). Normalized mutation densities per decile (y-axes) are presented for early (left) to late (right) replication domains. Normalized mutation densities of real somatic mutations and simulated somatic mutations from early to late replicating regions are shown as blue bars and dashed lines, respectively. ***(H)*** Enrichments and depletions of non-clustered, *omikli*, and *kataegis* mutations of B-cell malignancies within CTCF binding sites and histone modifications. Red colours correspond to enrichments of real somatic mutations when compared to simulated data. Blue colours correspond to depletions of real somatic mutations when compared to simulated data. The intensities of red and blue colours reflect the degree of enrichments or depletions based on the fold change. White colours correspond to lack of data for performing statistical comparisons. Statistically significant enrichments and depletions are annotated with * (q-value ≤ 0.05).

In summary, in this report we have performed a comprehensive topography analysis of mutational signatures encompassing 82,890,857 somatic mutations in 5,120 whole-genome sequenced tumours integrated with 516 tissue-matched topographical features from the ENCODE project. Our evaluation encompassed examining the effects of nucleosome occupancy, histone modifications, CTCF binding sites, replication timing, transcription strand asymmetry, and replication strand asymmetry on the accumulation of somatic mutations from more than 70 distinct mutational signatures. The results from these analyses have been provided as an online resource as a part of COSMIC signatures database, https://cancer.sanger.ac.uk/signatures/, where researchers can explore each mutational signature as well as each topographical feature in a cancer-specific manner.

## SUPPLEMENTARY FIGURE LEGENDS

**Figure S1. Somatic mutations in genic and intergenic regions imprinted by different mutational signatures. Related to Figure 1**. ***(A)*** Somatic mutations in genic and intergenic regions for signatures of single base substitutions (SBSs). Rows represent the signatures, where *n* reflects the number of cancer types in which each signature was found. Columns display the six substitution subtypes based on the mutated pyrimidine base: C>A, C>G, C>T, T>A, T>C, and T>G. SBS signatures with genic and intergenic regions asymmetries with q-values ≤ 0.05 are shown in circles with cyan and grey colours, respectively. The colour intensity reflects the odds ratio between the ratio of real mutations and the ratio of simulated mutations, where each ratio is calculated using the number of mutations in the genic regions and the number of mutations in the intergenic regions. Only odds ratios above 1.10 are shown. Circle sizes reflect the proportion of cancer types exhibiting a signature with specific genic versus intergenic regions asymmetry. ***(B)*** Somatic mutations in genic and intergenic regions for signatures of doublet-base substitutions (DBSs). Data are presented in a format similar to the one in panel *(A)*. ***(C)*** Somatic mutations in genic and intergenic regions for small insertions/deletions (IDs). Data are presented in a format similar to the one in panel *(A)*. ***(D)*** Histogram of fold enrichment as odds ratio between the ratio of real mutations and the ratio of simulated mutations, where each ratio is calculated using the number of mutations in the genic regions and the number of mutations in the intergenic regions. Frequency of fold enrichments (y-axis) are presented for discreet bins of fold enrichments (x-axis). Each fold enrichment reflects the odds ratio between real and simulated mutations where each ratio is the number of mutations in intergenic regions divided by the number of mutations in genic regions. Total number of fold enrichments, mean, and standard deviation of fold enrichments are shown in the upper right corner of the histogram. ***(E)*** Same format as panel *(D)* with the underlaying data reflecting fold enrichments after inflating the number of somatic mutations in genic regions to remove any transcription strand asymmetry.

**Figure S2. The effect of replication timing on mutational signatures. Related to Figure 2**. Top three panels reflect results for all single base substitutions (SBSs), all doublet-base substitutions (DBSs), and all small insertions/deletions (IDs) across all examined cancer types with each cancer type examined separately. Bottom panels reflect all somatic mutations attributed to a particular signature across all cancer types. Replication time data are separated into deciles, with each segment containing exactly 10% of the observed replication time signal (x-axes). Normalized mutation densities per decile (y-axes) are presented for early (left) to late (right) replication domains. Real data for SBS signatures are shown as blue bars, for DBS signatures as red bars, and for ID signatures as green bars. In all cases, simulated somatic mutations are shown as dashed lines. The total number of evaluated cancer types for a particular mutational signature is shown on top of each plot (*e.g.,* 36 cancer types were evaluated for SBS1). For each signature, the number of cancer types where the mutation density increases with replication timing is shown next to ⬈ (*e.g.,* 23 cancer types for SBS1). Similarly, the number of cancer types where the mutation density decreases with replication timing is shown next to ⬊ (*e.g.,* 0 cancer types for SBS1). Lastly, the number of cancer types where the mutation density is not affected by replication timing is shown next to ➞ (*e.g.,* 13 cancer types for SBS1).

**Figure S3. The effect of nucleosome occupancy on mutational signatures. Related to Figure 3**. Top three panels reflect results for all single base substitutions (SBSs), all doublet-base substitutions (DBSs), and all small insertions/deletions (IDs) across all examined cancer types with each cancer type examined separately. Bottom panels reflect all somatic mutations attributed to a particular signature across all cancer types. In all cases, solid lines correspond to real somatic mutations with blue solid lines reflecting SBSs, red solid lines reflecting DBSs, and green solid lines reflecting IDs. Solid lines and dashed lines display the average nucleosome signal (y-axes) along a 2 kilobase window (x-axes) centred at the mutation start site for real and simulated mutations, respectively. The mutation start site is annotated in the middle of each plot and denoted as 0. The 2 kilobase window encompasses 1,000 base-pairs 5’ adjacent to each mutation as well as 1,000 base-pairs 3’ adjacent to each mutation. For each mutational signatures, the total number of similar and considered cancer types using an *X/Y* format, with *X* being the number of cancer types where a signature has similar nucleosome behaviour (Pearson correlation ≥ 0.5 and q-value ≤ 0.05) and *Y* representing the total number of examined cancer types for that signature. For example, signature SBS3 annotated with 11/14 reflects a total of 14 examined cancer types with similar nucleosome behaviour observed in 11 of these 14 cancer types.

**Figure S4. Relationships between mutational signatures and histone modifications. Related to Figure 5**. *(A-C)* Relationships between 11 histone modifications and signatures of single base substitutions (SBSs) in panel *(A)*, doublet-base substitutions (DBSs) in panel *(B)*, and small insertions/deletions (IDs) in panel *(C)*. The examined histone modifications encompass H2AFZ, H3K4me1, H3K4me2, H3K4me3, H3K9ac, H3K9me3, H3K27ac, H3K27me3, H3K36me3, H3K79me2, and H4K20me1. Rows and columns reflect the mutational signatures and histone modifications, respectively. The circle in each cell is separated in red, blue, and grey segments proportionate to the cancer types in which the signature has a specific behaviour. A red segment in a circle reflects the signature being enriched in the vicinity of a histone modification (q-value ≤ 0.05 and at least 5% enrichment). A blue segment in a circle reflects the signature being depleted in the vicinity of a histone modification (q-value ≤ 0.05 and at least 5% depletion). A grey segment in a circle corresponds to neither depletion nor enrichment of the signature in the vicinity of a histone modification. Cells without a circle correspond to insufficient data to perform any statistical comparisons. ***(D)*** Exemplars of enrichment, depletions, or no effect for several histone modifications and signatures SBS7a and ID1. Solid lines and dashed lines display the average signal for a particular histone modification (y-axes) along a 2 kilobase window (x-axes) centred at the mutation start site for real and simulated mutations, respectively. The mutation start site is annotated in the middle of each plot and denoted as 0. The 2 kilobase window encompasses 1,000 base-pairs 5’ adjacent to each mutation as well as 1,000 base-pairs 3’ adjacent to each mutation.

**Figure S5. Strand asymmetries of non-clustered, *omikli*, and *kataegis* substitutions across 288 whole-genome sequenced B-cell malignancies. Related to Figure 7**. Transcription strand asymmetries are shown in the left panels where bars display the six substitution subtypes based on the mutated pyrimidine base: C>A, C>G, C>T, T>A, T>C, and T>G (depicted on the x-axes). Y-axes correspond to the numbers of single base substitutions. Blue bars reflect real transcribed substitutions, while shaded blue bars correspond to simulated transcribed substitutions. Similarly, green bars reflect real untranscribed mutations, whereas shaded green bars correspond to simulated untranscribed substitutions. Replication strand asymmetries are shown in the middle panels where bars display the six substitution subtypes based on the mutated pyrimidine base: C>A, C>G, C>T, T>A, T>C, and T>G (depicted on the x-axes). Y-axes correspond to the numbers of single base substitutions. Red bars reflect real substitutions on the lagging strand, while shaded red bars correspond to simulated substitutions on the lagging strand. Similarly, yellow bars reflect real substitutions on the leading strand, whereas shaded yellow bars correspond to simulated substitutions on the leading strand. Comparisons of genic and intergenic regions are shown in the right panels where bars display the six substitution subtypes based on the mutated pyrimidine base: C>A, C>G, C>T, T>A, T>C, and T>G (depicted on the x-axes). Y-axes correspond to the numbers of single base substitutions. Cyan bars reflect real substitutions in genic regions, while shaded cyan bars correspond to simulated substitutions in genic regions. Similarly, grey bars reflect real substitutions in intergenic regions, whereas shaded grey bars correspond to simulated substitutions in intergenic regions. Results for non-clustered mutations are shown in panel *(A)*, *omikli* mutations in panel *(B)*, and *kataegis* mutations in panel *(C)*. Statistically significant strand asymmetries are shown with stars: * q-value ≤ 0.05; ** q-value ≤ 0.01; *** q-value ≤ 0.001.

## SUPPLEMENTARY TABLES

**Supplementary Table 1: External Datasets Utilized in Performing the Topography Analyses.** The table provides information on the ENCODE (https://www.encodeproject.org/) or GEO (https://www.ncbi.nlm.nih.gov/gds) datasets used for examining every topography feature within each evaluated cancer type.

## Supporting information

STAR Methods

## ACKNOWLEDGEMENTS

The authors would like to thank the COSMIC team for assistance in developing and deploying the COSMIC signatures topography database. BO and LBA would like to thank Prof. Steven Rozen (Duke-NUS) and Prof. Michael Stratton (Sanger Institute) for the many useful discussion in regard to examining the topography of mutational signatures as well as Mariya Kazachkova (UC San Diego) for her feedback and comments on improving the readability of the methods. This was work was funded by Cancer Research UK Grand Challenge Award [C98/A24032] as well as US National Institute of Health grants R01ES030993, R01ES032547, and R01CA269919. Work at the Wellcome Sanger Institute was also supported by the Wellcome Trust [grant number 108413/A/15/D]. LBA is also supported by a Packard Fellowship for Science and Engineering. The funders had no roles in study design, data collection and analysis, decision to publish, or preparation of the manuscript.

## AUTHOR CONTRIBUTIONS

BO and LBA conceived the performed computational analyses and wrote the manuscript with assistance from MDG. BO developed the Python code and performed the bioinformatics analyses with assistance from MDG, ENB, MZ, and MB. The online COSMIC signatures topography database was designed by BO, IV, and LBA with assistance from MDG, MZ, and MB. The COSMIC signatures topography database was implemented by IV with feedback from all authors. LBA supervised the overall development of the code, website, analysis, and writing of the manuscript. All authors read and approved the final manuscript.

