## Supplementary material for "Topography of mutational signatures in human cancer": Figure S1

Figure S1. Somatic mutations in genic and intergenic regions imprinted by different mutational signatures. Related to Figure 1.

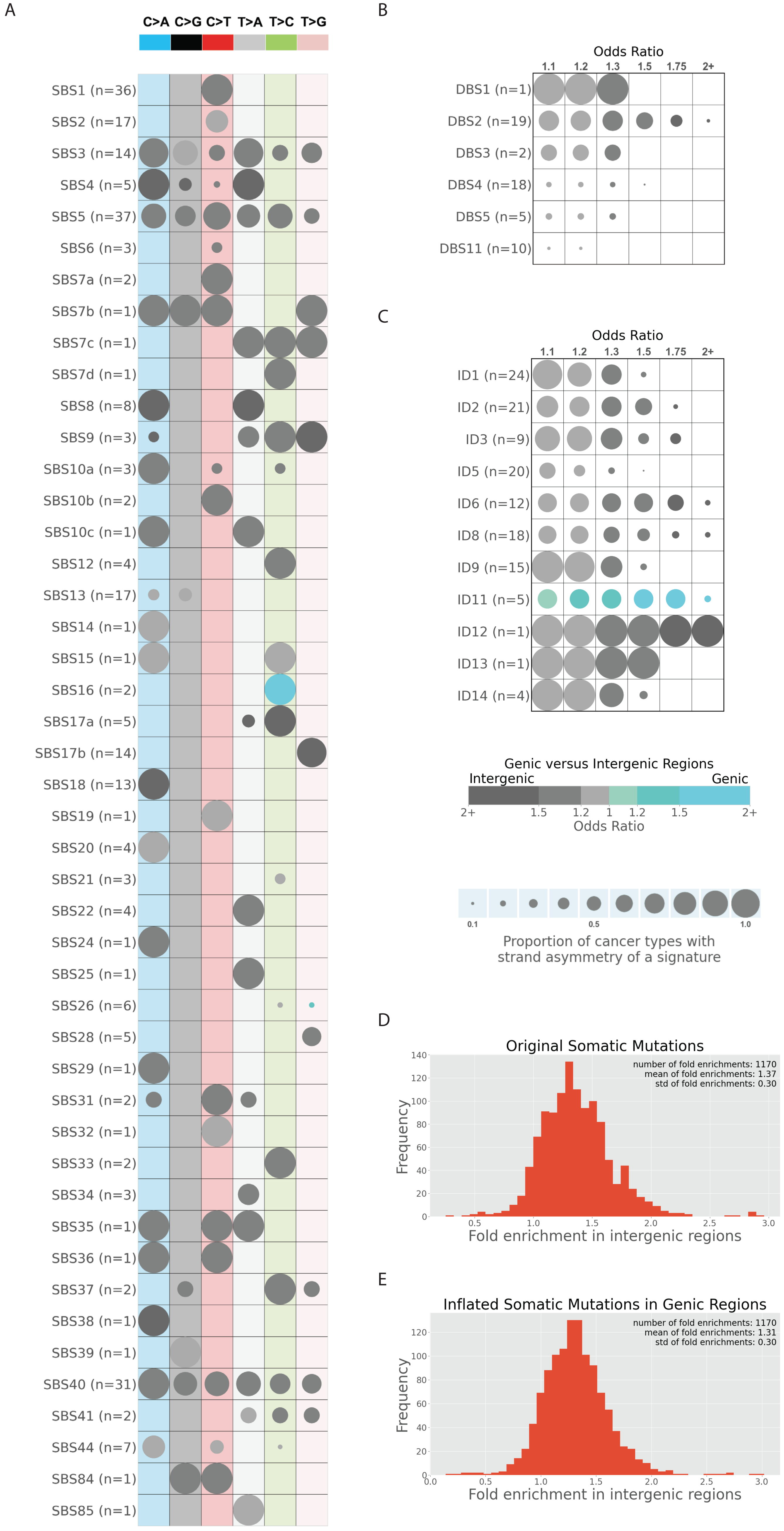
