## Supplementary figures and images for "Topography of mutational signatures in human cancer"

### Figure S2

Figure S2. The effect of replication timing on mutational signatures. Related to Figure 2.

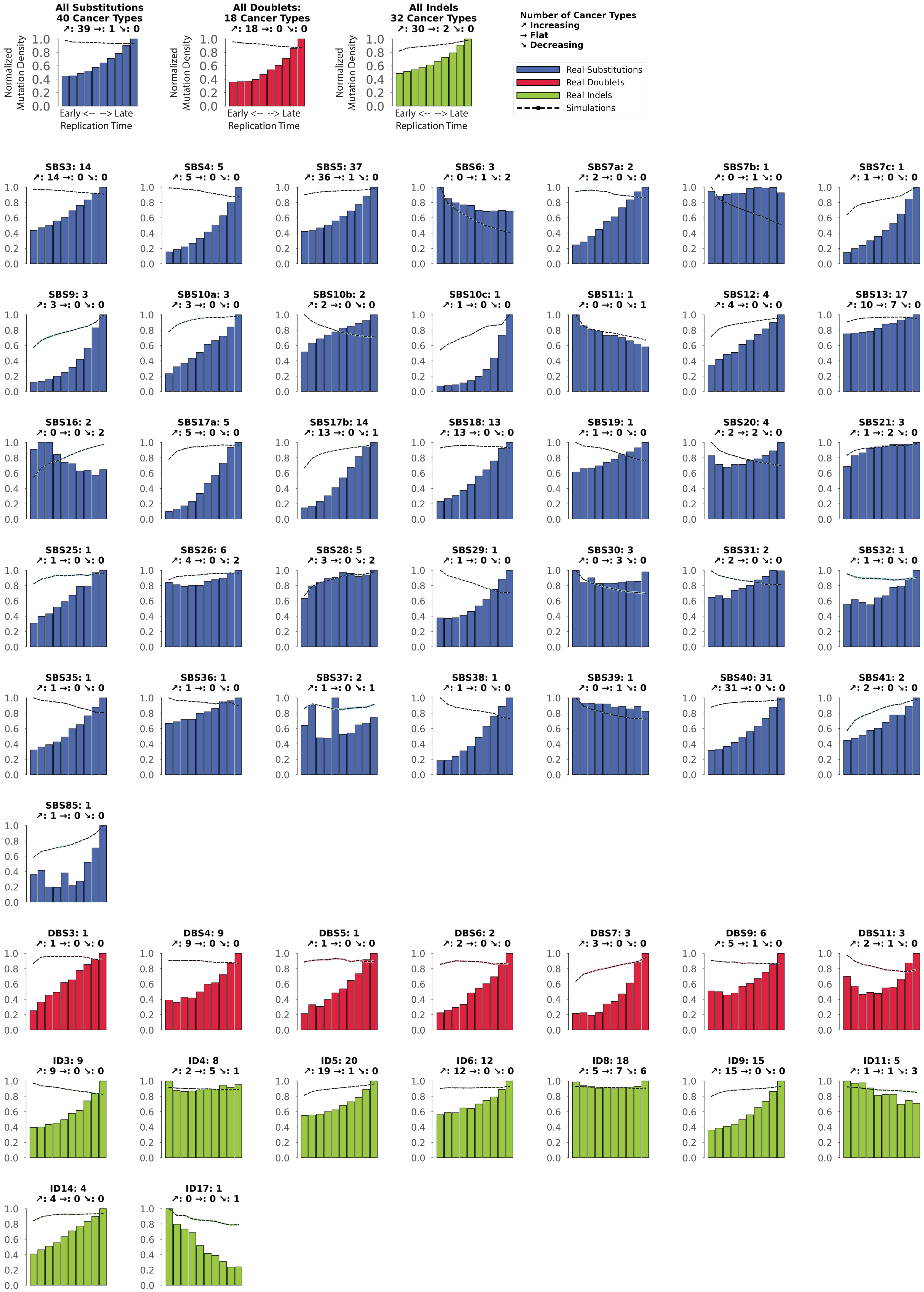

### Figure S3

Figure S3. The effect of nucleosome occupancy on mutational signatures. Related to Figure 3.

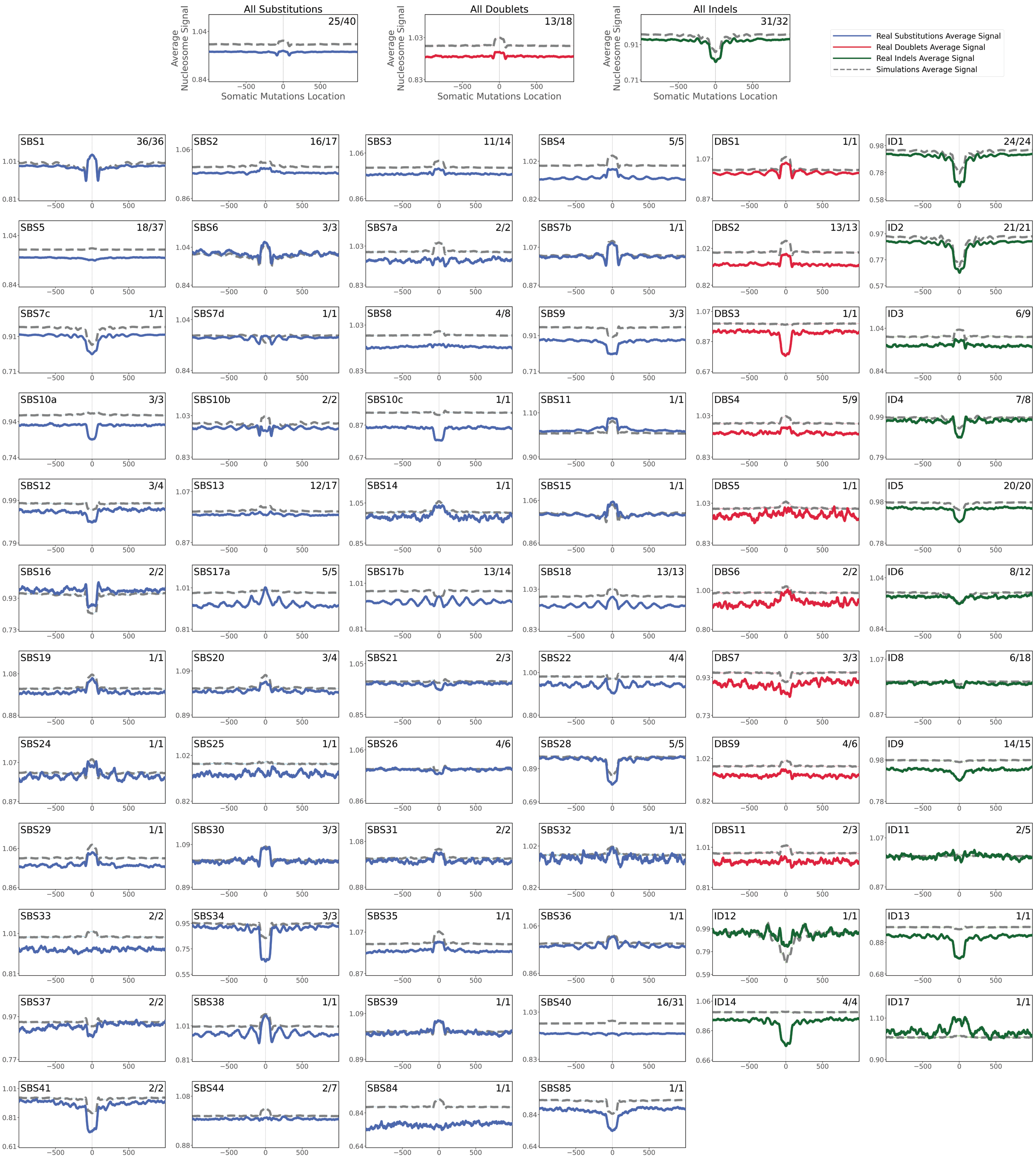
