## Supplementary material for "Topography of mutational signatures in human cancer": Figure S4

Figure S4. Relationships between mutational signatures and histone modifications. Related to Figure 5.

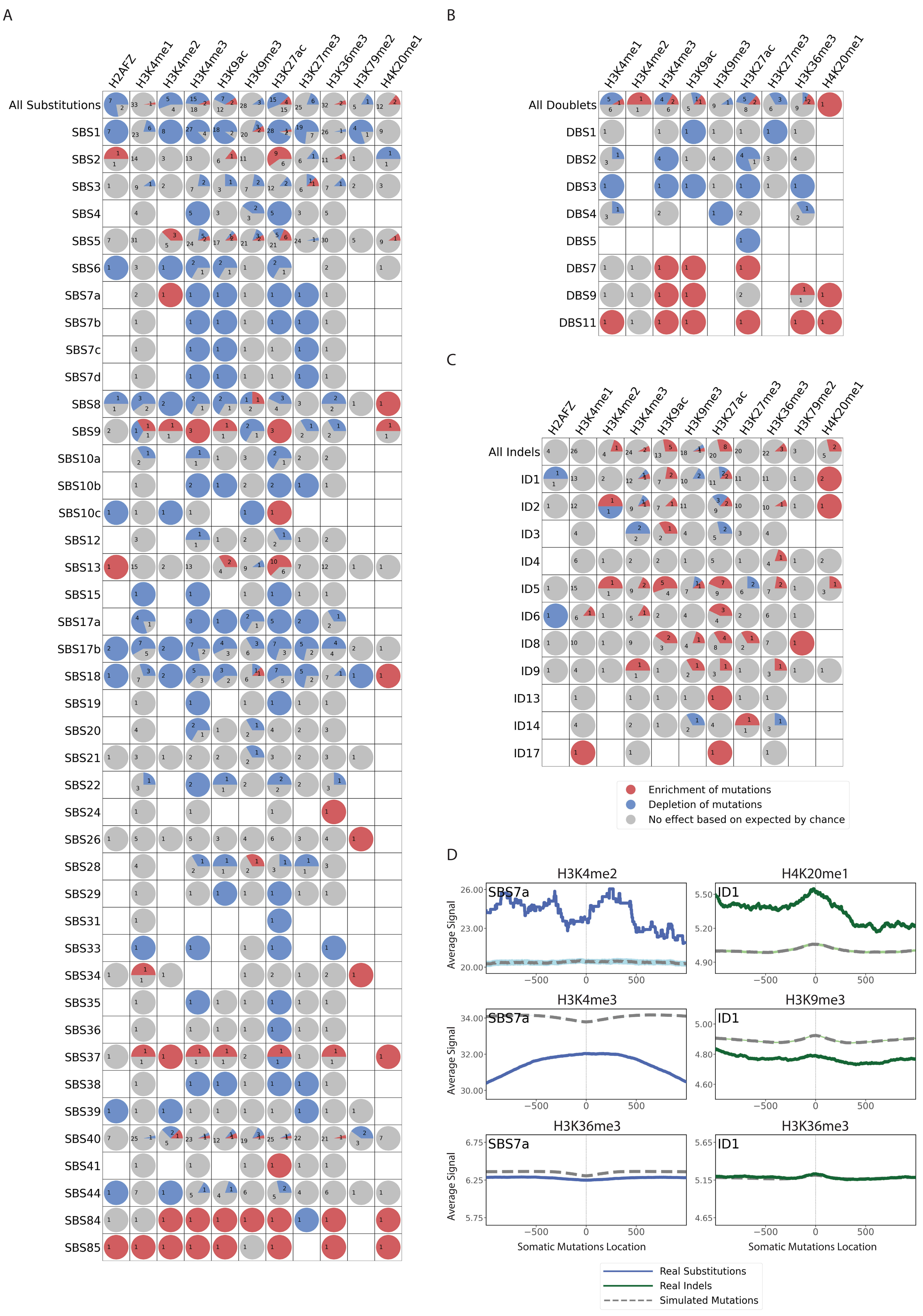
