## Supplementary material for "Topography of mutational signatures in human cancer": Figure S5

Figure S5. Strand asymmetries of non-clustered, omikli, and kataegis substitutions across 288 whole-genome sequenced B-cell malignancies. Related to Figure 7.

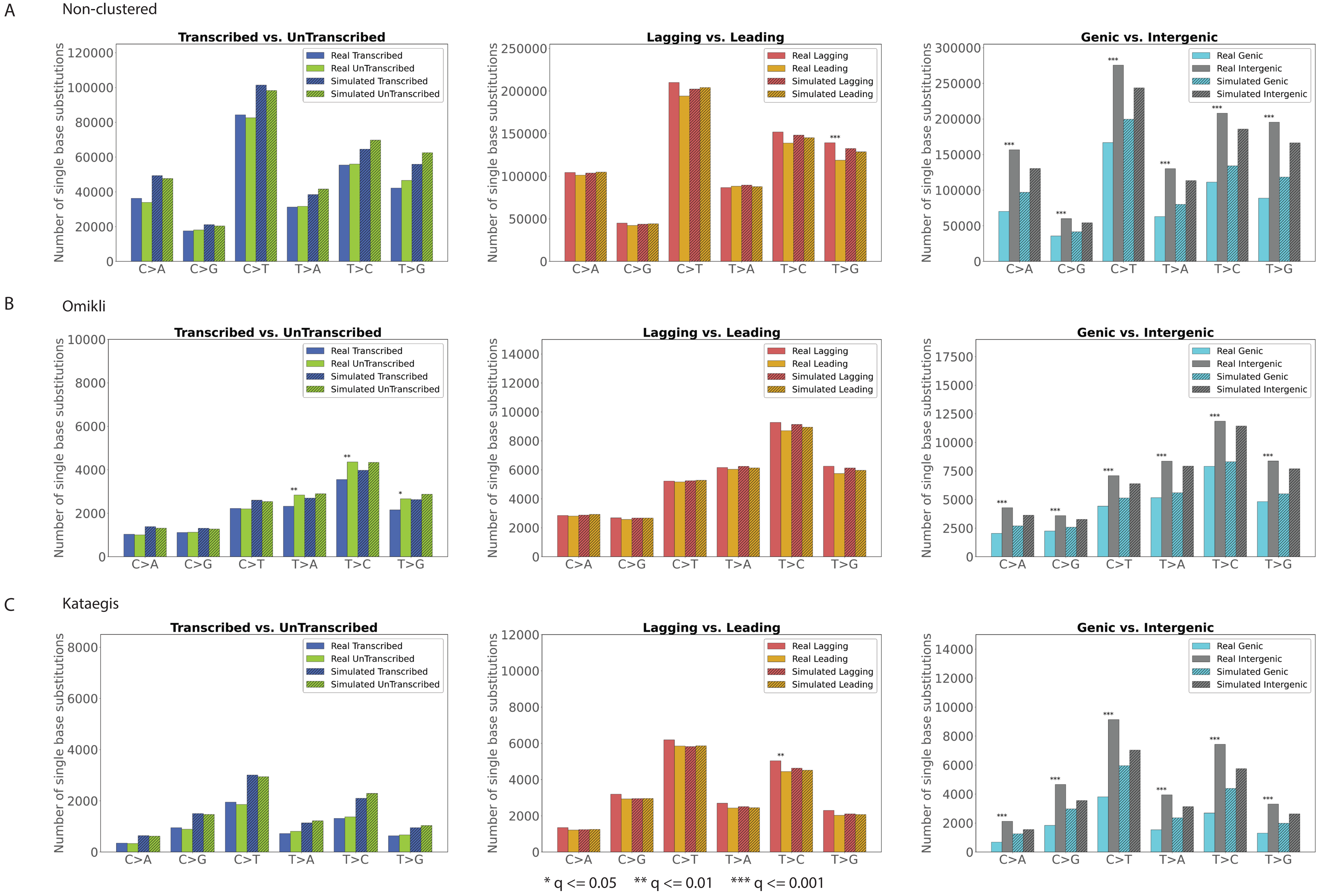
