## Supplementary material for "Topography of mutational signatures in human cancer": STAR Methods

### KEY RESOURCES TABLE

| REAGENT or RESOURCE | SOURCE | IDENTIFIER |
|---|---|---|
| **Biological samples** | | |
| PCAWG project (core WGS dataset)<br>Somatic Mutations and Mutational Catalogues from PCAWG Project | Alexandrov et al.[1] | https://www.synapse.org/#!Synapse:syn11804058<br>https://www.synapse.org/#!Synapse:syn11804040 |
| PCAWG project (additional WGS dataset)<br>VCF like sample files and SBS signatures in samples | Alexandrov et al.[1] | https://www.synapse.org/#!Synapse:syn11801872<br>https://www.synapse.org/#!Synapse:syn11801496 |
| MUTOGRAPHS project | Moody et al.[2] | https://doi.org/10.6084/m9.figshare.22744733 |
| **Deposited data** | | |
| Topography of Mutational Signatures in Human Cancer | This paper | COSMIC Signatures v3.3<br>https://cancer.sanger.ac.uk/signatures |
| **Software and algorithms** | | |
| SigProfilerMatrixGenerator (v1.1.31) | Bergstrom et al.[3] | https://github.com/AlexandrovLab/SigProfilerMatrixGenerator |
| SigProfilerSimulator (v1.1.2) | Bergstrom et al.[4] | https://github.com/AlexandrovLab/SigProfilerSimulator |
| SigProfilerExtractor (v1.1.0) | Islam et al.[5] | https://github.com/AlexandrovLab/SigProfilerExtractor |
| SigProfilerClusters (v1.0.11) | Bergstrom et al.[6] | https://github.com/AlexandrovLab/SigProfilerClusters |
| SigProfilerTopography (v1.0.70) | This paper | https://github.com/AlexandrovLab/SigProfilerTopography |
| Cancer-type specific and across all cancer-types combined topography analysis | This paper | https://github.com/AlexandrovLab/SigProfilerTopographyCombined |
| bigWigToWig tool (v446 April 2023) | Kent et al.[7] | http://hgdownload.cse.ucsc.edu/admin/exe/ |

| | | |
|---|---|---|
| liftOver tool (v446 April 2023) | Kent et al.[7] | http://hgdownload.cse.ucsc.edu/admin/exe/ |
| Python (v3.7.0) | Python Software Foundation | https://www.python.org/ |
| Python package: pandas (v1.1.5) | McKinney[8] | https://pandas.pydata.org |
| Python package: NumPy (v1.20.1) | Harris et al.[9] | https://numpy.org |
| Python package: matplotlib (v3.4.2) | Hunter[10] | https://matplotlib.org |
| Python package: SciPy (v1.6.3) | Virtanen et al.[11] | https://scipy.org |
| Python package: statsmodels (v0.12.2) | Seabold and Perktold[12] | https://www.statsmodels.org |
| **Other** | | |
| Transcription factors (TF) binding sites datasets (TF ChIP-seq assays) | ENCODE Project | htts://www.encodeproject.org/ Exact file names for each utilized dataset are available as part of Supplementary Table 1 |
| Histone modifications sites datasets (Histone ChIP-seq assays) | ENCODE Project | https://www.encodeproject.org/ Exact file names for each utilized dataset are available as part of Supplementary Table 1 |
| Nucleosome occupancy datasets (MNase-seq assays) | ENCODE Project | https://www.encodeproject.org/ Exact file names for each utilized dataset are available as part of Supplementary Table 1 |
| Replication timing datasets (Repli-seq assays) | ENCODE Project | https://www.encodeproject.org/ Exact file names for each utilized dataset are available as part of Supplementary Table 1 |

### RESOURCE AVAILABILITY

#### Lead contact

#### Materials availability

This study did not generate new unique reagents beyond the analysed data and the developed source code (see below).

#### Data and code availability

- All topographical data and figures regarding topography of mutational signatures in human cancer generated in this study have been deposited at COSMIC, Catalogue of Somatic Mutations in Cancer (https://cancer.sanger.ac.uk/signatures/), through COSMIC Signatures v3.3, released on May 27th, 2022 and are currently publicly available.

- All original Python code has been deposited on GitHub and is publicly available as of the date of publication. Links to GitHub repositories are listed in the Key Resources Table.

- This paper analyses existing publicly available datasets. Accession numbers for the datasets are listed in the Key Resources Table.

- Any additional information required to reanalyse the data reported in this paper is available from the lead contact upon request.

### METHODS DETAILS

#### Simulating Synthetic Cancer Datasets

Synthetic cancer datasets were simulated using SigProfilerSimulator[4]. Briefly, the tool randomly generated single base substitutions (SBSs), doublet base substitutions (DBSs), and small insertions and deletions (IDs) while maintaining the patterns of the original somatic mutations in each sample at a preselected resolution. Simulations were performed 100 times for each examined cancer genome while maintaining the mutational burden on each chromosome in each sample. All simulations were performed using SBS-96, DBS-78, and ID-83 mutational classification schemas[3]. Briefly, SBS-6 represents single base substitutions in 6 mutational classes (C>A, C>G, C>T, T>A, T>C, T>G) considering the pyrimidine base of the Watson-Crick base-pair for each somatic mutation. SBS-96 is a further expansion of SBS-6 mutational classification by adding the immediate 5' and 3' adjacent bases for each somatic mutation within the representation of every mutation. DBS-78 catalogues doublet-base substitutions in 78 mutational channels using the maximum pyrimidine context of the Watson-Crick base-pairs[3], whereas ID-83 classifies small insertions and deletions into 83 mutational channels by considering the size of the ID event and the repeat size surrounding the insertion or deletion event[3].

#### Assigning Signature Probabilities to Somatic Mutations

The performed topography analyses are based on the assignment of signature probabilities to each individual somatic mutation. For this purpose, SigProfilerExtractor was utilized for *de novo* extraction of mutational signatures and decomposition of *de novo* extracted signatures to the set of reference COSMIC mutational signatures[1,5]. After Poisson resampling and normalization of the original mutational matrix for each replicate, SigProfilerExtractor performs nonnegative matrix factorization for multiple iterations to identify an optimal solution. Briefly, SigProfilerExtractor identifies the optional decomposition rank *k* by performing decompositions with different ranks and applying consensus clustering to identify a stable solution that best explains the underlying data[1,5]. After extracting *de novo* mutational signatures, each *de novo* signature is matched to a COSMIC mutational signature and the COSMIC signatures are assigned using a penalized nonnegative least square approach[1,5]. Moreover, SigProfilerExtractor automatically assigns a probability for each operative signature to generate every individual mutation within all examined samples[1,5].

**Matching Cancer Types with ENCODE Datasets**

Experimental data were downloaded from ENCODE for each evaluated topographical feature (**Table S1**). When multiple ENCODE datasets were used for the same topographical feature in a cancer type, analyses were performed for all ENCODE datasets and the results were averaged across the examined datasets. Any ENCODE genomic coordinates reported using GRCh38 annotations were first remapped to GRCh37 annotations using liftOver with exclusion of any ambiguously mapping regions[7]. ENCODE files in bigWig file format were converted into wig files using bigWigToWig file format conversion software[7].

We analysed a total of 82,890,857 somatic mutations (79,269,539 single base substitutions, 429,179 doublet-base substitutions, and 3,192,139 small insertions and deletions) from 40 cancer types derived using 5,120 whole-genome sequenced samples from PCAWG, PCAWG other, and MUTOGRAPHS projects[1,2]. Cancer types were matched to the closest available ENCODE datasets (**Table S1**). Topography analyses were performed both across all cancer types as well as within each individual cancer-type. The global pan-cancer analyses are shown in the manuscript while all individual cancer-type analyses are available through the COSMIC database: https://cancer.sanger.ac.uk/signatures/

**Annotating somatic mutations based on cellular transcription**

Somatic mutations were called with respect to + strand of the reference genome and annotated in regard to the pyrimidine base of the mutated base pair[3]. Specifically, SigProfilerMatrixGenerator[3] was used for examining transcriptional strand asymmetry for single base substitutions, doublet base substitutions, and small indels. The tool evaluates whether a mutation occurs on the transcribed or the non-transcribed strand of well-annotated protein coding genes of the human reference genome. Mutations found in the transcribed regions of the human genome are further subclassified as *transcribed* or *un-transcribed*. Any mutations in bi-directionally transcribed regions were ignored in the current analysis. Additionally, mutation found outside the transcribed regions of the human genome are subclassified as *non-transcribed*. In all cases, mutations were oriented based on the reference strand and their pyrimidine context.

**Annotating somatic mutations based on cellular replication**

For each cancer type of interest, genomic regions were annotated either as being on the leading or being on the lagging strand using our previously developed approach[13]. Briefly, analyses were performed for tissue-matched wavelet-smoothed replication timing signal data incorporated with valleys (replication termination zones) and peaks (replication initiation zones) data. Valleys and peaks were sorted with respect to their genomic coordinate in ascending order. Each consecutive stretches of DNA of at least 10 kilobases long with positive slope corresponded to leading strand regions on the positive strand, whereas negative slope provided lagging strand regions on the positive strand. We discarded the latest 25 kilobases of the replication termination zones to be stringent in our annotations. Having annotated genome regions as leading regions (+ slope) and lagging regions (– slope) on the positive strand, we automatically acquired leading regions (- slope) and lagging regions (+ slope) on the negative strand. Mutations were counted as being on leading strand or lagging strand based on their occupancy in a leading or lagging region. Similar to the annotation for transcription, in all cases, mutations were first oriented based on the reference strand and their pyrimidine context.

**Detecting strand asymmetries across cancer types**

For each mutational signature and for all cancer types having this mutational signature, we retrieved the number of mutations on each strand/region in six mutational channels (C>A, C>G, C>T, T>A, T>C, and T>G). P-values were calculated for the odds ratio between the ratio of real mutations and the ratio of simulated mutations. Specifically, for transcription strand asymmetry odds ratios were calculated between the ratios of real mutations and the ratios of simulated mutations, where each ratio is calculated using the number of mutations on the transcribed strand and the number of mutations on the untranscribed strand. Similarly, for replication strand asymmetry odds ratios were calculated between the ratios of real mutations and the ratios of simulated mutations, where each ratio is calculated using the number of mutations on the lagging strand and the number of mutations on the leading strand. Lastly, for genic and intergenic regions, odds ratios were calculated between the ratios of real mutations and the ratios of simulated mutations, where each ratio is calculated using the number of mutations in the genic regions and the number of mutations in the intergenic regions The computed p-values were corrected for multiple testing using Benjamini-Hochberg method. Only strand asymmetries with corrected p-value ≤ 0.05 and odds ratios above 1.10 were considered and reported as part of the presented results.

**Detecting strand-coordinated mutagenesis of mutational signatures**

Analyses of strand-coordinated mutagenesis searched for consecutive single base substitutions on the same DNA strand with inter-mutational distance less than 10,000 base-pairs within the same sample as previously done for breast cancer in[13]. To find the consecutive mutations, all the single base substitutions in a sample were first assigned to the SBS signature with the highest probability, and only the mutations with the probability greater than or equal to a pre-set cut-off value of at least 0.50 were retained (*i.e.*, at least 50% chance for a signature to have generated that mutation). Somatic mutations were sorted in an ascending order in regard to chromosomal positions and consecutive groups of substitutions with the same mutational context, on the same DNA strand, and attributed to the same substitution signature were identified. Where applicable, consecutive groups of substitutions were combined with the appropriate adjustments of their group lengths. Any consecutive groups of substitutions with length of 1 were discarded. All results coming across different samples were pooled. For each SBS signature and strand-coordinated mutagenesis group length, the observed number of groups for real mutations and the expected number of groups coming from 100 simulated datasets were compared with the simulated datasets serving as null hypotheses. P-values were computed using z-tests evaluating whether the mean values of the expected number of groups were equal to the mean values of the observed number of groups for each simulated dataset. The computed p-values were corrected for multiple testing using

Benjamini-Hochberg method. SBS signatures and strand-coordinated mutagenesis group lengths with corrected p-value $\leq 0.05$ were considered and reported as part of the presented results.

**Analyses of replication timing**

As previously done in breast cancer, wavelet-smoothed signal data were used in the replication timing analysis[13]. Briefly, cancer types were matched with ENCODE data from the most suitable tissue or cell line and the corresponding Repli-seq dataset was utilized in the analysis (**Table S1**). Given the Repli-Seq signal data for a tissue of interest, a higher replication time signal reflects an earlier replication timing. The replication time signals were each sorted in a descending order and, subsequently, the sorted replication time signals were divided into deciles. Each decile contains approximately 10% of each replication time signal. Somatic mutations of interest were distributed within the corresponding deciles based on their overlap with the replication domains in the examined deciles. To correct for genomic size, mutation densities were calculated by dividing the numbers of somatic mutations within each decile by the number of attributable bases of adenine, thymine, guanine, and cytosine (excluding any ambiguous genomic annotations in the reference genome). To compare replication timing of different mutational signatures with each other, mutation densities were further normalized with respect to the highest mutation density observed for each respective signature. Lastly, as with other analyses, the reported replication timing analyses included only signatures with at least 1,000 somatic mutations unambiguously attributed to an individual mutational signature.

**Mutational signature and cancer type specific replication timing analysis**

To compare the replication timing between real and simulated somatic mutations, cancer-type specific normalized mutation densities were calculated for real mutations, $x_{real} = [x_{real}^{1}, x_{real}^{2}, \dots, x_{real}^{10}]^{T}$, and for each of the 100 simulated synthetic cancer datasets, $X_{sim\_i} = [x_{sim\_i}^{1}, x_{sim\_i}^{2}, \dots, x_{sim\_i}^{10}]^{T}$, where $sim\_i = 1, 2, \dots, 100$. Normalized mutation density vectors generated based on each simulated dataset were combined and the matrix $\boldsymbol{X_{sims}} = [X_{sim\_1}, X_{sim\_2}, \dots, X_{sim\_100}]^{T}$ was generated. Mean simulated vector, $\overline{x}_{sims}$, standard deviation vector, $\sigma_{sims}^{X}$, and their 95% confidence intervals were calculated using SciPy. As a result, normalized mutation densities of real mutations across replication timing deciles, $x_{real}$, were compared to the averaged normalized mutation densities of simulated mutations across replication timing deciles, $\overline{x}_{sims}$. Results were further averaged across all cancer types in order to generate the summary plots presented in the manuscript.

To classify whether the mutation density was increasing, flat, or decreasing in regards to replication timing, we fitted a linear regression model to the values of the normalized mutation densities, $x_{real}$. A mutational signature was considered to be increasing from early to late replicating regions if the slope $m$ was statistically significant from a flat line and the values of $x_{real}$ were monotonically increasing. A mutational signature was considered to be decreasing from early to late replicating regions if the slope $m$ was statistically significant from a flat line and the values of $x_{real}$ were monotonically decreasing. Lastly, a mutational signature was considered to be generally unaffected by replication timing if the slope $m$ was not statistically significant from a flat line.

**Occupancy analysis of topographical features**

Genomic occupancy analysis evaluated the relationship between mutational signatures and the genomic locations of different topographical features, including: *(i)* nucleosomes; *(ii)* transcription factors; and *(iii)* histone modifications. Specifically, occupancy analysis of topographical features focused on mutations within a specific genomic window, and it evaluated the average experimental signal of a particular topographical feature. In all cases, this window was centred on a somatic mutation, and the window began 1,000 base-pair (bp) 5' of a mutation and ended 1,000 bp 3' of a mutation; for example, this resulted in a region 2,001 bp for each examined somatic single base substitution. For a given topographical feature, the analysis evaluated the experimental signal for a set of cancer-matched datasets from ENCODE by averaging the signal across the regions of interest. For example, to evaluate the connection between SBS2 and CTCF binding in breast adenocarcinoma, our analysis utilized 4 datasets of chromatin immunoprecipitation followed by sequencing (ChIP–seq) from breast tissues in ENCODE (**Table S1**); for each mutation unequivocally attributed to SBS2 in the examined breast cancers, the CTCF signals within 2,001 bp around the mutation were averaged across the 4 examined datasets. As in other analysis, the reported occupancy results included only signatures with at least 1,000 unequivocally attributed somatic mutations to that specific mutational signature within each examined cancer type.

Next, we averaged both the real and simulated mutations in two rounds where the first round of accumulation and averaging was across all mutations for each cancer-type matched ENCODE dataset and the second round of accumulation and averaging was across all cancer-type matched ENCODE datasets. For real mutations, in the first round, for each cancer-type matched ENCODE dataset, we accumulated the average signal vectors coming from all real somatic mutations. This resulted in a cancer-type specific vector $K_{real} = [k_{real}^0, k_{real}^1, \dots, k_{real}^{2000}]^T$, where $K_{real}$ is the average signal of the topographical feature of interest in a 2,001 bp window using all real mutations. In the second round, we accumulated the average signal vectors $K_{real}$ that were attained in the first round coming from each cancer-type matching ENCODE dataset and derived their average for the total number of considered ENCODE datasets. This results in a global vector $M_{real}$ $= [m_{real}^0, m_{real}^1, \dots, m_{real}^{2000}]^T$, where $M_{real}$ represents the average signal of the topographical feature in a 2,001 bp window using all cancer-type matching ENCODE datasets. The same procedure was repeated for each of the 100 simulated synthetic cancer datasets resulting in an average $K_{sims}$ for a signature and topographical feature within each cancer type as well as an average $M_{sims}$ for a signature and topographical feature across all cancer types. For each signature and topographical feature, comparisons between $K_{real}$ and $K_{sims}$ were performed within each cancer type while a global comparison was performed between $M_{real}$ and $M_{sims}$. Linear correlations were performed to evaluate whether the occupancy signal within ± 500 base-pair windows around a somatic mutation for a cancer type, $K_{real}$, is correlated with the average signal across all cancer types $M_{real}$. Any statistically significant Pearson's correlations, based on Benjamini-Hochberg corrected p-value ≤ 0.05, were reported as part of the presented results.

**Abundance analysis of mutational signatures and topographical features**

In addition to performing occupancy analysis, our abundance analysis evaluated the enrichment, depletion, or no relationship between mutational signatures and topography features of interest. Specifically, we investigated the relationships between mutational signatures and the following topographical features: *(i)* nucleosomes; *(ii)* transcription factors; and *(iii)* histone modifications. For each mutational signature and topography feature, we examined whether there is an enrichment, depletion, or no statistically significant relation between the mutational signature and the topography feature of interest by comparing the real somatic mutations with the sets of simulated somatic mutations. Specifically, the average signal vector of real mutations for each cancer-type matched ENCODE dataset, $K_{real}$, was obtained as described in the occupancy analysis. An average value, $s_{real}$, was derived for ± 50 bp window centred at the somatic mutations for each cancer-type matched ENCODE dataset. Similar analysis was performed for the 100 simulated cancer datasets, which allowed deriving a $K_{sim}$ average vector and $s_{sim}$ average value.

To evaluate whether this average signal value of real mutations, $s_{real}$, was expected by chance given the average signal values coming from 100 simulations, $s_{sim\_i}$ for $i = 1, 2, \dots, 100$, we assessed the statistical significance of each fold change and associated it with a p-value. Average signal value for real mutations was determined as observed value, $s_{real}$, and average signal values coming from 100 simulations were determined as expected values, $\left[ s_{sim\_1}, s_{sim\_2}, \dots, s_{sim\_100} \right]^T$. Z-test was applied to test whether the observed value was the mean of expected values under null hypothesis and a p-value together with a test statistic were obtained. The computed p-values were corrected for multiple testing using Benjamini-Hochberg method and only p-values ≤ 0.05 were considered and reported as part of the presented results. In case of multiple ENCODE datasets availability for a certain cancer type and topography feature of interest, the calculated p-values coming from each cancer-type matched ENCODE dataset were pooled and combined using Fisher's method. In this cases, p-values were corrected for multiple testing using Benjamini-Hochberg method after combining. Likewise, calculated fold changes acquired from each ENCODE dataset were averaged and average fold change was obtained.

**Classifying cancers into ones with low and ones with high APOBEC3 mutations**

Each cancer sample with operative APOBEC3-associated signatures SBS2 or SBS13 was classified as either having low, mid, or high APOBEC3 presence. Specifically, we utilized a previously developed scheme for calculating APOBEC3 presence[14], where a ratio was derived as the natural logarithm of the total number of mutations attributed to the APOBEC3-associated signatures SBS2 or SBS13 divided by the natural logarithm of the total number of mutations excluding ones due to APOBEC3-associated signatures SBS2 or SBS13. Thus, for each sample the ratio was calculated in the following manner:

$$\text{ratio} = \frac{\log_e(\text{number of mutations attributed to SBS2 or SBS13})}{\log_e(\text{total number of mutations NOT attributed to SBS2 and SBS13})}$$

Cancers with ratios ≥0.90 were classified as samples with high APOBEC3 presence and cancers with ratios ≤0.75 were classified as samples with low APOBEC3 presence. All samples with ratios between 0.75 and 0.90 were classified as mid APOBEC3 presence and were not considered in our subsequent replication timing re-analysis. Specifically, we repeated the replication timing for all cancer types by separately examining samples with low APOBEC3 presence and separately examining samples with high APOBEC3 presence. As done in the prior analysis in this manuscript, all results were averaged within and across the examined cancer types.

**QUANTIFICATION AND STATISTICAL ANALYSIS**

All statistical analysis were performed in Python using NumPy, SciPy, statsmodels, and other standard statistical modules. Statistical parameters and details of the analyses can be found in "Method Details".

### ADDITIONAL RESOURCES

All topographical data and figures regarding topography of mutational signatures in human cancer generated in this study were deposited at COSMIC, Catalogue of Somatic Mutations in Cancer (https://cancer.sanger.ac.uk/signatures/), through COSMICv3.3, May 27th, 2022.
